# Development and validation of serological markers for detecting recent exposure to *Plasmodium vivax* infection

**DOI:** 10.1101/481168

**Authors:** Rhea J. Longley, Michael T. White, Eizo Takashima, Jessica Brewster, Masayuki Morita, Matthias Harbers, Leanne J. Robinson, Fumie Matsuura, Shih-Jung Zoe Liu, Connie S. N. Li-Wai-Suen, Wai-Hong Tham, Julie Healer, Christele Huon, Chetan E. Chitnis, Wang Nguitragool, Wuelton Monteiro, Carla Proietti, Denise L. Doolan, Xavier C. Ding, Iveth J. Gonzalez, James Kazura, Marcus Lacerda, Jetsumon Sattabongkot, Takafumi Tsuboi, Ivo Mueller

**Affiliations:** Population Health and Immunity Division, Walter and Eliza Hall Institute of Medical Research, Melbourne, Australia; Department of Medical Biology, University of Melbourne, Melbourne, Australia.; Mahidol Vivax Research Unit, Faculty of Tropical Medicine, Mahidol University, Bangkok, Thailand.; Malaria Parasites & Hosts Unit, Department of Parasites & Insect Vectors, Institut Pasteur, Paris, France.; Division of Malaria Research, Proteo-Science Center, Ehime University, Matsuyama, Japan.; CellFree Sciences Co., Ltd., Yokohama, Japan.; Burnet Institute, Melbourne, Australia.; Infection and Immunity Division, Walter and Eliza Hall Institute of Medical Research, Melbourne, Australia.; Malaria Parasite Biology and Vaccines, Department of Parasites & Insect Vectors, Institut Pasteur, Paris, France.; Department of Molecular Tropical Medicine and Genetics, Faculty of Tropical Medicine, Mahidol University, Bangkok, Thailand.; Fundacão de Medicina Tropical Dr. Heitor Vieira Dourado, Manaus, Brazil.; Universidade do Estado do Amazonas, Manaus, Brazil.; Centre for Molecular Therapeutics, Australian Institute of Tropical Health and Medicine, James Cook University, Cairns, Australia.; QIMR Berghofer Medical Research Institute, Brisbane, Australia.; Foundation for Innovative New Diagnostics, Geneva, Switzerland.; Center for Global Health and Diseases, Case Western Reserve University, Cleveland, United States.; Instituto Leônicas & Maria Deane (Fiocruz), Manaus, Brazil.; RIKEN Center for Integrated Medical Sciences (IMS), Yokohama, Japan.

**Author notes:** These authors contributed equally (Rhea J. Longley, Michael T. White).

**Keywords:** malaria, *Plasmodium vivax*, serological markers, exposure, surveillance, naturally acquired immunity, hypnozoites, malaria elimination, RBP2b, RAMA, MSP1, MSP3, MSP8, EBP, Pv-fam-a

## Abstract

In order to accelerate towards malaria elimination, improved targeting of limited resources is essential. A major gap in our elimination toolkit for *Plasmodium vivax* malaria is the identification of individuals carrying arrested liver stages, called hypnozoites. These clinically silent but frequently relapsing hypnozoites are key to *P. vivax* persistence. Whilst hypnozoites cannot be directly detected, individuals who have had recent exposure to *P. vivax* and have not been treated are likely to harbor these parasites. By measuring IgG antibody responses to over 300 *P. vivax* proteins, a panel of serological markers capable of detecting exposure to *P. vivax* infections in the prior 9-month period was identified and validated. Using antibody responses to 8 *P. vivax* proteins, 80% sensitivity and specificity for detecting recent infections were achieved in three independent studies conducted in Thailand, Brazil and the Solomon Islands. As these individuals have a high likelihood of harboring hypnozoites, the suite of these 8 antibody responses can serve as biomarkers for the identification of individuals who should be targeted for treatment with liver-stage drugs such as primaquine and tafenoquine in mass drug administration programs aimed at controlling and eliminating *P. vivax* malaria.

**One Sentence Summary:** The manuscript describes identification and validation of a novel panel of P. vivax proteins that can be used to detect recent exposure to P. vivax infections within the prior 9 months.

## Introduction

Elimination of malaria by the year 2030 is now the explicit goal of all malaria endemic countries in the Asia-Pacific. While impressive progress towards this goal has been made, global funding for malaria control has flat-lined since 2010 and progress has stalled in many parts of the world (*1*). New interventions and tools for better targeting of limited resources are urgently needed.

A major hurdle for elimination is the increasing proportion of malaria cases caused by *Plasmodium vivax* as transmission declines (*1, 2*). *P. vivax* has a number of unique biological features that make its control difficult (*3*), including very high prevalence of low density, asymptomatic infections (*4*) and an arrested liver-stage that can reactivate weeks to months later to cause relapsing infections that cause morbidity as well as sustain transmission. These hypnozoites, which are not detectable by any current diagnostic method, can be responsible for over 80% of all blood-stage infections (*5*). Identifying and targeting individuals with hypnozoites is thus essential for accelerating and achieving malaria elimination. In addition, as endemicity decreases, malaria transmission becomes increasingly fragmented and highly seasonal (*6, 7*) rendering blanket approaches to malaria control and elimination inefficient. This requires new, innovative tools and approaches specifically designed to assist in targeting interventions to the changing malaria epidemiological landscape (*8*).

National malaria control programs still rely almost exclusively on microscopy or rapid diagnostic tests for the routine detection of malaria cases at health clinics and for surveillance using mass blood surveys or reactive case-detection (RACD) (*9*). While relatively cheap and readily available, these tools have limited sensitivity for detecting individuals with asymptomatic malaria who have lower parasite densities (*10*), making it difficult for control programs to efficiently identify and delineate areas of low and high *P. vivax* transmission and target their resources accordingly. Molecular techniques such as PCR have much greater sensitivity (*10*) but are rarely implemented by control programs due to high cost and the need for a specialised laboratory. All of these methods can only detect individuals with a current blood-stage infection, rendering mass screening and treatment (MSAT) approaches ineffective for reducing *P. vivax* transmission because they fail to treat individuals that only carry hidden *P. vivax* liver-stage infections (*5, 11*) with no circulating blood-stage parasitemia. Mass drug administration (MDA) is predicted to be highly effective as a control tool but only if it also includes a drug that targets *P. vivax* hypnozoites (*5*). Unfortunately, primaquine and tafenoquine, the only current drugs able to eliminate hypnozoites, can have toxic side effects in glucose-6-phosphate dehydrogenase (G6PD) deficient individuals (*12*), limiting their acceptability for MDA campaigns without prior G6PD screening, in particular in low transmission areas where >90% MDA recipients will have no direct benefit from the treatment. There is thus a diagnostic gap, with MSAT ineffective due to under treatment and MDA unacceptable due to overtreatment. A strategy that can efficiently target individuals with current blood-stage infections and those with hypnozoites is required.

Blood-stage *P. vivax* infections induce a robust antibody response in humans to a broad range of *P. vivax* proteins, even following low-density asymptomatic infections (*13*). IgG antibody responses to specific *Plasmodium* proteins can be long-lived even after clearance of blood-stage infection (*14, 15*), and hence antibodies are markers of past exposure as well as current infections (*15*). Population-level serological exposure markers (SEM) have been used for surveillance of malaria and a number of other infectious diseases (i.e. leishmaniasis, influenza, trachoma), and antibody responses can be simply and cheaply measured in point-of-care tests (*16-18*). Besides general surveillance, SEMs may also be used for risk stratification and as guidance for targeted malaria control and elimination interventions (*19*). For *P. vivax* there is an important additional, individual-level application for SEMs: to target treatment to people at-risk of carrying clinically silent hypnozoites. While it is not possible to directly detect hypnozoites with current technology, all tropical and sub-tropical *P. vivax* strains cause a primary infection followed by a first relapse after no more than 6-9 months (*20-22*). Therefore, any individual with blood-stage infection within the previous 9 months, who has not received anti-liver-stage drugs, is likely to be a hypnozoite carrier. A carefully selected panel of proteins inducing antibodies that signify exposure within the previous 9 months could provide an accurate measure of current transmission and also highlight individuals likely to harbour hypnozoites who could be targeted for treatment with liver-stage drugs, such as primaquine or tafenoquine, after serological testing (“serological testing and treatment”, seroTAT).

342 *P. vivax* proteins in this study, identified from the *P. vivax* genome (*23*), were screened for their ability to detect recent exposure to the parasite, and their use was subsequently validated in malaria-endemic regions of Thailand, Brazil and the Solomon Islands.

## Results

### Antigen discovery phase

Data on the magnitude and longevity of IgG responses to 342 *P. vivax* proteins following symptomatic *P. vivax* infections (*24*) were used to identify suitable proteins for detecting recent exposure (Fig. 1a). IgG was measured in four longitudinal plasma samples from individuals in Thailand (n=32) and Brazil (n=33) to enable estimation of antibody half-life (*24*) (Table S1).

**Fig. 1.**
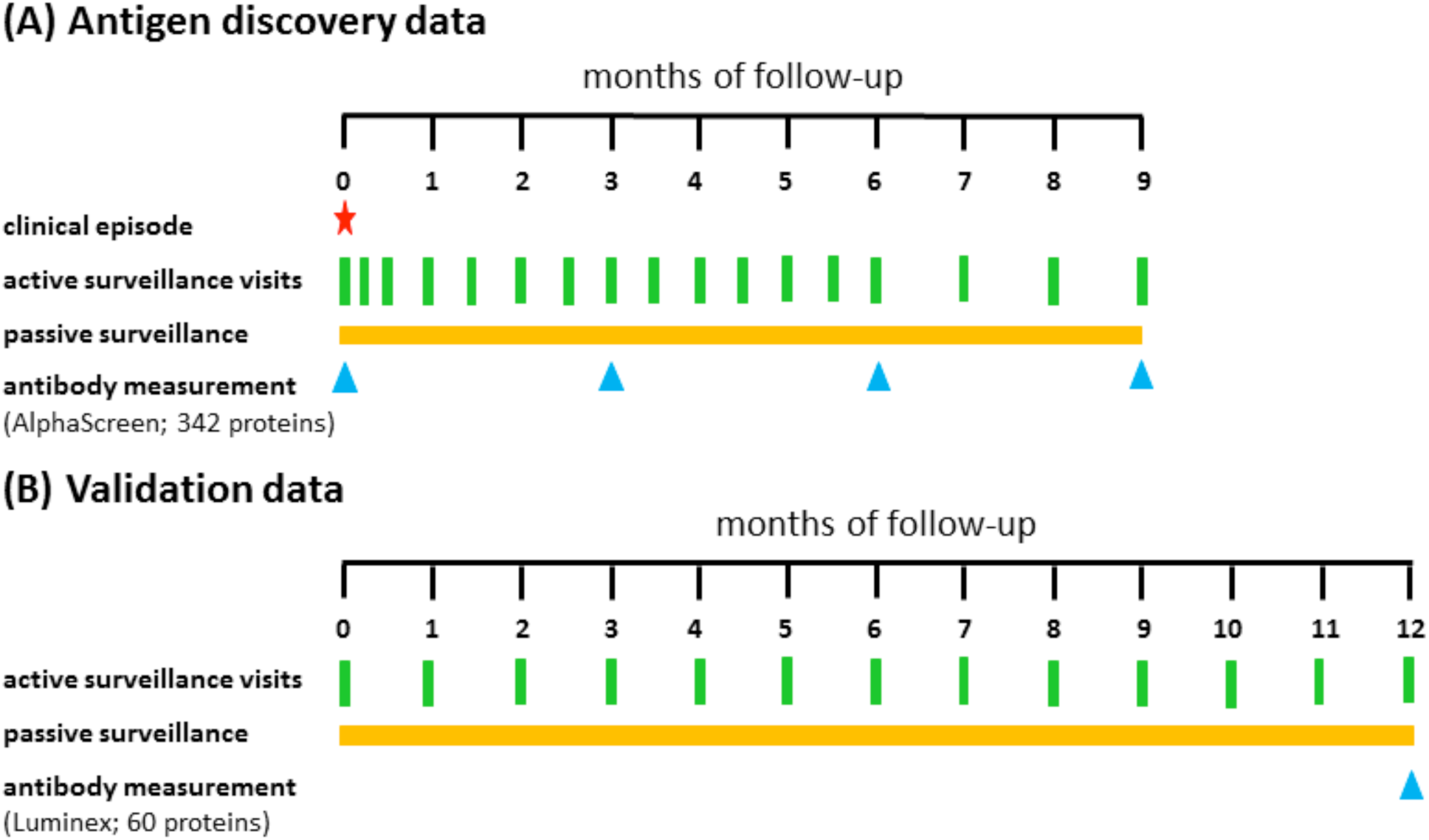
Study design and follow-up schedule. (A) Thai and Brazilian patients were enrolled following a clinical episode of *P. vivax* and treated according to the relevant National Guidelines, with treatment directly observed (DOT) to ensure compliance and reduced risk of relapse. Volunteers were followed for 9 months after enrolment, with finger-prick blood samples collected at enrolment and week 1, then every 2 weeks for 6 months, then every month. Antibody levels were measured in a subset of 32 Thai and 33 Brazilian volunteers who were confirmed to be free of blood-stage *Plasmodium* parasites by analysing all samples by light microscopy and qPCR. (B) 999 participants from Thailand, 1274 participants from Brazil, and 860 participants from the Solomon Islands were followed longitudinally for 12 months with active surveillance visits every month. Antibody levels were measured in plasma samples from the last visit.

A down-selection pipeline was developed to identify candidate serological markers using this data (Fig. 2). *P. vivax* proteins were first prioritised, with selection of those that had similar estimated IgG half-lives in both antigen discovery cohorts (Thailand and Brazil), and those that were highly immunogenic at the time of infection (at least 50% seropositive individuals) (Fig. 2a). Using these, 142 of the 342 *P. vivax* proteins were prioritised as having suitable characteristics for use as candidate SEMs.

**Fig. 2.**
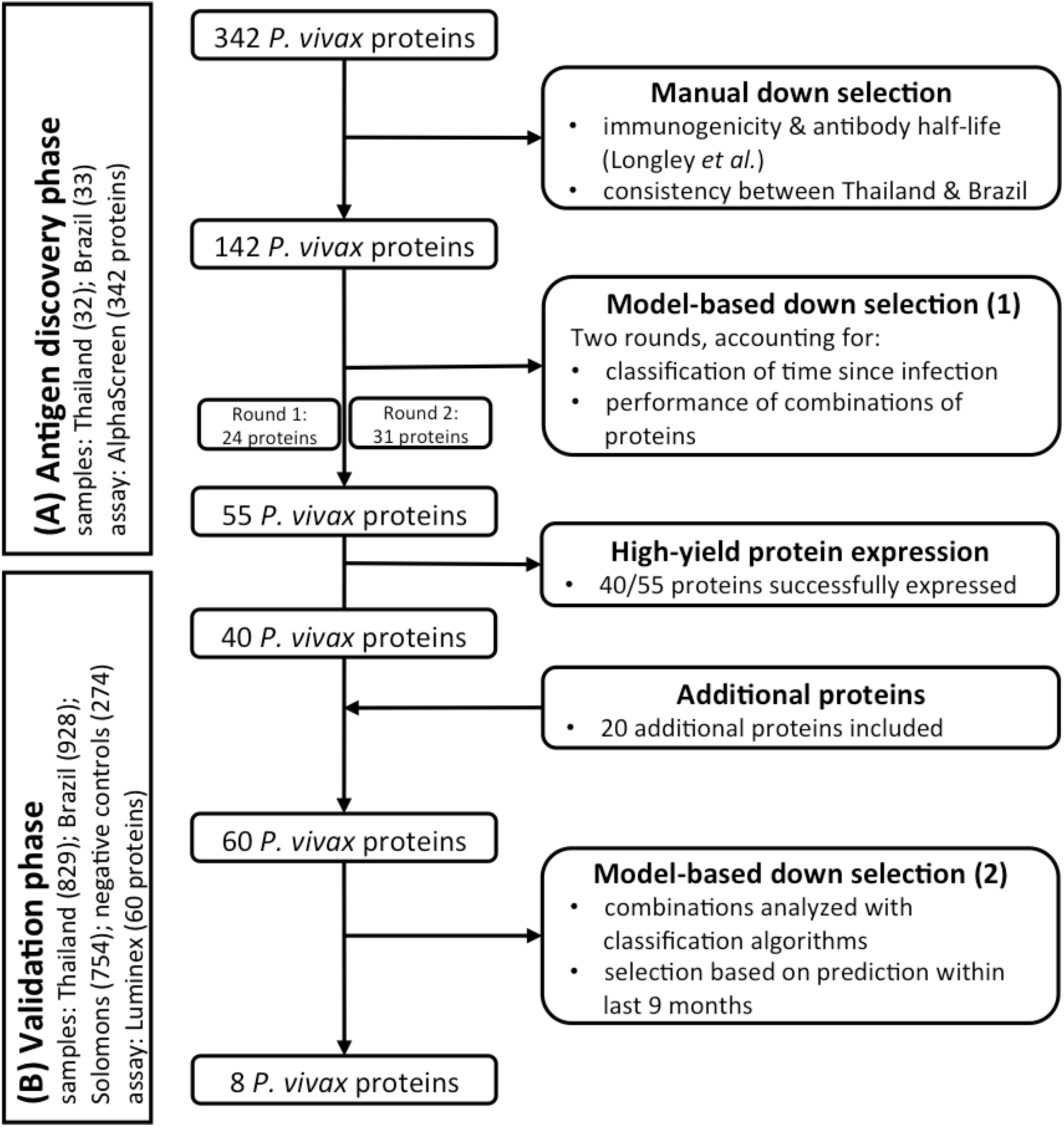
Flow diagram of antigen down-selection pipeline. The studies and samples used are listed to the left in (A) and (B), whereas the pipeline and number of proteins included at each stage are listed to the right.

A statistical model for estimating time since infection was used to test the ability of the 142 prioritised proteins, individually or in combination, to predict time since last infection. Fig. 3a,b shows an example of dynamics of antibody responses to five proteins in two representative individuals from Thailand and Brazil. Fig. 3c,d shows the estimated time since last infection with uncertainty for antibody responses measured to the 5 proteins in Fig. 3a,b at 6 months after infection, for the same two representative individuals. Increasing the number of proteins to at least five resulted in higher accuracy compared to using a single protein alone, with diminishing returns beyond 20 proteins (Fig. S4). A simulated annealing algorithm was employed to determine optimal combinations of proteins that maximise the likelihood of correctly estimating the time since last infection (Fig. 3e). Whilst some proteins had 100% probability of being included in a successful panel of a set size (Fig. 3e), there was some redundancy in choice of additional proteins and hence all were ranked based on their probability of inclusion in a 20-protein panel (given the diminishing returns beyond 20 proteins). The 24 highest ranked proteins were selected for further testing.

**Fig. 3.**
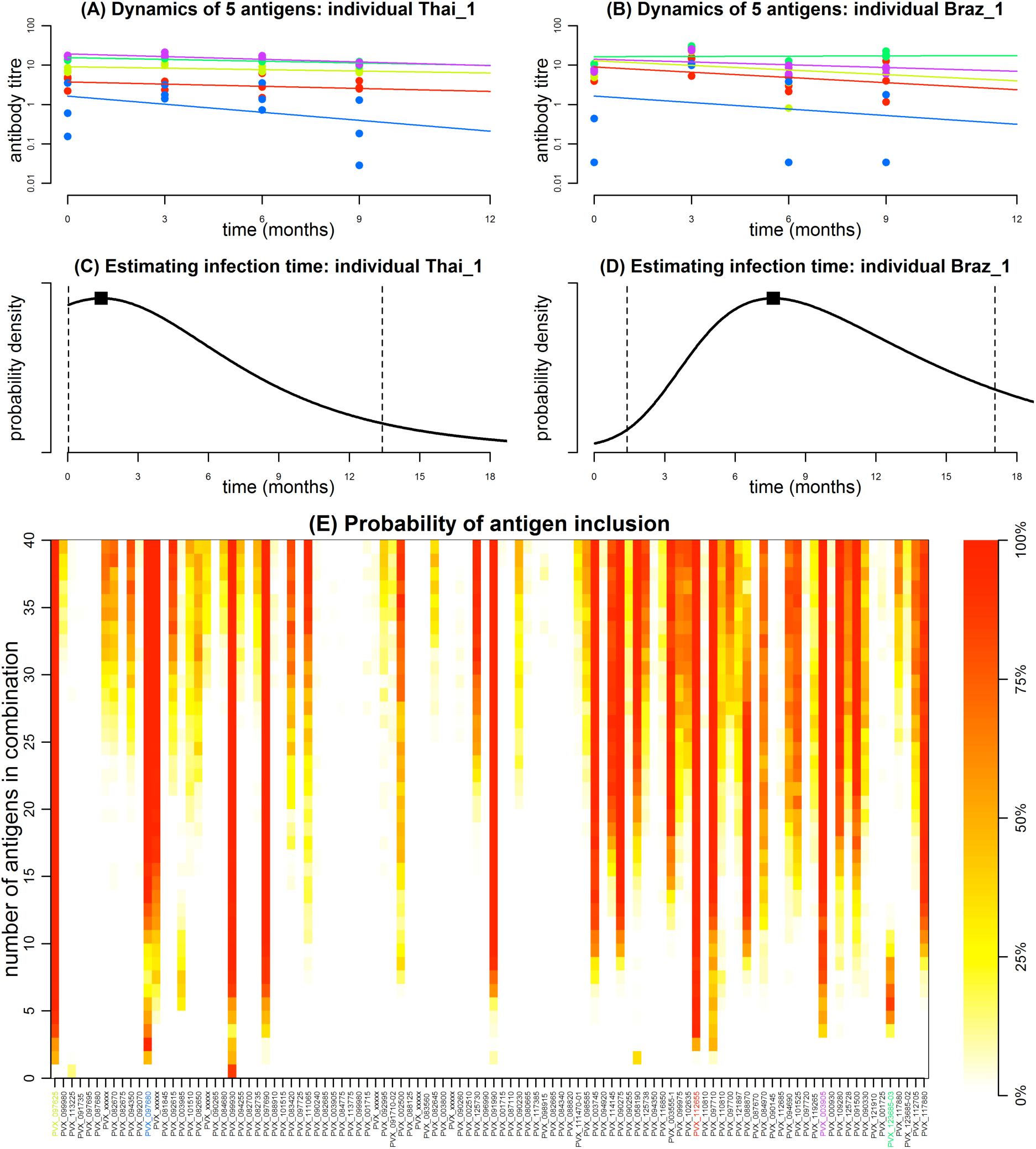
Antigen discovery phase: estimating time since last *P. vivax* infection. (A, B) Examples of the dynamics of antibody responses to five proteins in individuals from Thailand and Brazil. Points represent measured antibody responses in triplicate on the AlphaScreen™ platform, and solid lines depict the fit of mixed-effects linear regression models. Each colour corresponds to one of the five proteins (PVX_097625, PVX_097680, PVX_112655, PVX_003905, PVX_123685-03) selected to provide optimal classification performance. (C, D) Using data on measured antibody responses 6 months after infection, the black curve shows the model estimated time since infection using the antibody responses shown in (A, B). The square point denotes the most likely estimate and dashed vertical lines denote the 95% confidence interval. In both cases the confidence intervals span 6 months. (E) Results of simulated annealing algorithm for estimating the probability that combinations of antigens predict time since last infection in the antigen discovery data sets. Colour is representative of the inclusion probability (red 100%, white 0%). The five proteins with data shown in figures a-d are highlighted in colour in panel e. These results came from the second round of down-selection on 104 *P. vivax* proteins (see Supplementary Methods for further details).

The wheat germ cell free (WGCF) system was selected for expression of the proteins due to its eukaryotic nature and high rates of past success in expressing *Plasmodium* proteins (*25*). Despite all proteins being expressed in this system previously as crude proteins (*24*) not all could be produced at high yield and high purity, and thus an additional round of down-selection using ranking from the simulated annealing algorithm was performed to increase the number of proteins retained at this point in the pipeline. The algorithm was run on 104 of the 142 prioritised proteins, excluding the top 24 already selected (to provide more discriminatory power for selecting additional proteins) and 14 proteins known to have low yields or aggregates from our previous work. An additional 31 proteins were selected in the second round using the same methodology, with 55 *P. vivax* proteins in total identified as suitable candidate SEMs. Of these, 40 were successfully produced at a high yield and purity using the WGCF system (Table 2, Table S2).

**Table 2.** Purified *P. vivax* proteins used in the validation phase and their individual performance. Area under the curve (AUC) from the single antigen classifier is shown; proteins are listed in order of best individual performance. 8 of 60 proteins are shown here; see Table S2 for the extended and complete Table. References listed are for the protein production and purification method.

| Protein ID<sup>a</sup> | Gene Annotation<sup>a</sup> | Protein length, aa | Construct, aa (size) | Expression System<sup>c</sup> | Purification Method | AUC |
| --- | --- | --- | --- | --- | --- | --- |
| PVX_094255B | reticulocyte binding protein 2b (RBP2b) | 2806 | 161-1454 (1294) | *E. coli* | 2x affinity + size exclusion (27, 30, 57) | 0.819 |
| PVX_099980 | merozoite surface protein 1 (MSP1-19) | 1751 | 1622-1729 (108) | WGCF | One-step Ni column | 0.804 |
| PVX_000930 | sexual stage antigen s16, putative | 140 | 31-end (110) | WGCF | One-step Ni column | 0.78 |
| PVX_094255A | reticulocyte binding protein 2b (RBP2b) | 2806 | 1986-2653 (667) | WGCF | One-step Ni column | 0.779 |
| PVX_087885B | rhoptry-associated membrane antigen, putative | 730 | 462-730 (269) | WGCF | One-step Ni column (28) | 0.774 |
| PVX_095055 | Rh5 interacting protein, putative (RIPR) | 1075 | 552-1075 (524) | *E. coli* | 2x affinity + size exclusion (59) | 0.767 |
| AAY34130.1<sup>b</sup> | Duffy binding protein (DBP, region 2, AH strain) | 237 | 1-237 (237) | *E. coli* | Ni, ion exchange, gel filtration (58) | 0.764 |
| PVX_097715 | hypothetical protein | 450 | 20-end (431) | WGCF | One-step Ni column | 0.76 |

<sup>a</sup>PlasmoDB release 36 (http://plasmodb.org/plasmo/), <sup>b</sup>GenBank

An additional 20 proteins that were known to be highly immunogenic were added to these 40 proteins (*26-29*). This included four proteins already within the down-selected panel of 40 (different constructs or produced by a different laboratory and protein expression systems), and three distinct constructs of PVX_110810 (Duffy binding protein, each representing a different region of the protein or sequence from a different strain) (Table 2, Table S2).

### Validation phase

The next step in the down-selection pipeline was to validate the use of the 60 candidate SEMs in larger geographically diverse cohorts with a known history of malaria infections in the preceding 12 months, using plasma samples from three, one-year long observational cohort studies conducted in Thailand, Brazil and the Solomon Islands (Fig. 1b, Table 1). Individuals in these cohorts were assessed monthly for the presence of *Plasmodium spp.* by qPCR, with continuous concurrent passive case detection at local malaria clinics and hospitals. Plasma samples from the last visit of these cohorts (n=829, 928 and 754 in Thailand, Brazil and the Solomon Islands, respectively) were used to measure IgG responses in a multiplex Luminex® assay to the 60 *P. vivax* proteins. 158/2511 individuals in these cohorts had a concurrent *P. vivax* infection, detected by qPCR, at the time the plasma was collected; IgG antibody levels were strongly associated with current infection status for each of the 60 proteins in the Thai cohort (OR 2·1-7·7, p<0·05) and for the majority of proteins in the Brazilian cohort (57/60, OR 1·5-7·1, p<0·05) (Table S3). This association was not as clear for the paediatric Solomon Islands cohort, with IgG levels to only 29 out of 60 proteins significantly associated with current *P. vivax* infections (OR 1·7-6·3, p<0·05). Overall, there was a pattern of decreasing IgG magnitude with increasing time since last *P. vivax* infection (Fig. 4a-h, Fig. S5), with minimal reactivity in malaria naïve negative control individuals from Bangkok and Melbourne.

**Table 1.** Epidemiological overview of data sets. Individuals in the antigen discovery phase were recruited following a clinical episode of *P. vivax*. *denotes PCR prevalence in the general population.

| population | N: number of participants | n: subset with antibody measurements | age (years) | sex | PCR P<sub>v</sub>PR | time since last PCR positive sample |  |  |  |
| --- | --- | --- | --- | --- | --- | --- | --- | --- | --- |
|  |  |  | median (range) | % female |  | current | <9 months | 9-12 months | no detected |
| (A) Antigen discovery data |  |  |  |  |  |  |  |  |  |
| Thailand | 57 | 32 | 29 (7, 71) | 44% | 11.4%* | — | — | — | — |
| Brazil | 91 | 33 | 36 (16, 56) | 21% | 5.0%* | — | — | — | — |
| (B) Validation data |  |  |  |  |  |  |  |  |  |
| Thailand | 999 | 829 | 24 (1, 78) | 54.7% | 3.33% | 25 | 47 | 25 | 732 |
| Brazil | 1274 | 928 | 25 (0, 103) | 50.7% | 5.21% | 40 | 165 | 31 | 692 |
| Solomon Islands | 860 | 754 | 5.6 (0.5, 12.7) | 51.6% | 11.92% | 93 | 172 | 29 | 566 |
| Victorian Blood Donor Registry (VBDR) | 102 | 102 | 39 (18, 68) | 69.6% | 0.0% | 0 | 0 | 0 | 102 |
| Australian Red Cross (ARC) | 100 | 100 | 52 (18, 77) | 54.0% | 0.0% | 0 | 0 | 0 | 100 |
| Thai Red Cross (TRC) | 72 | 72 | — | — | 0.0% | 0 | 0 | 0 | 72 |

**Fig. 4.**
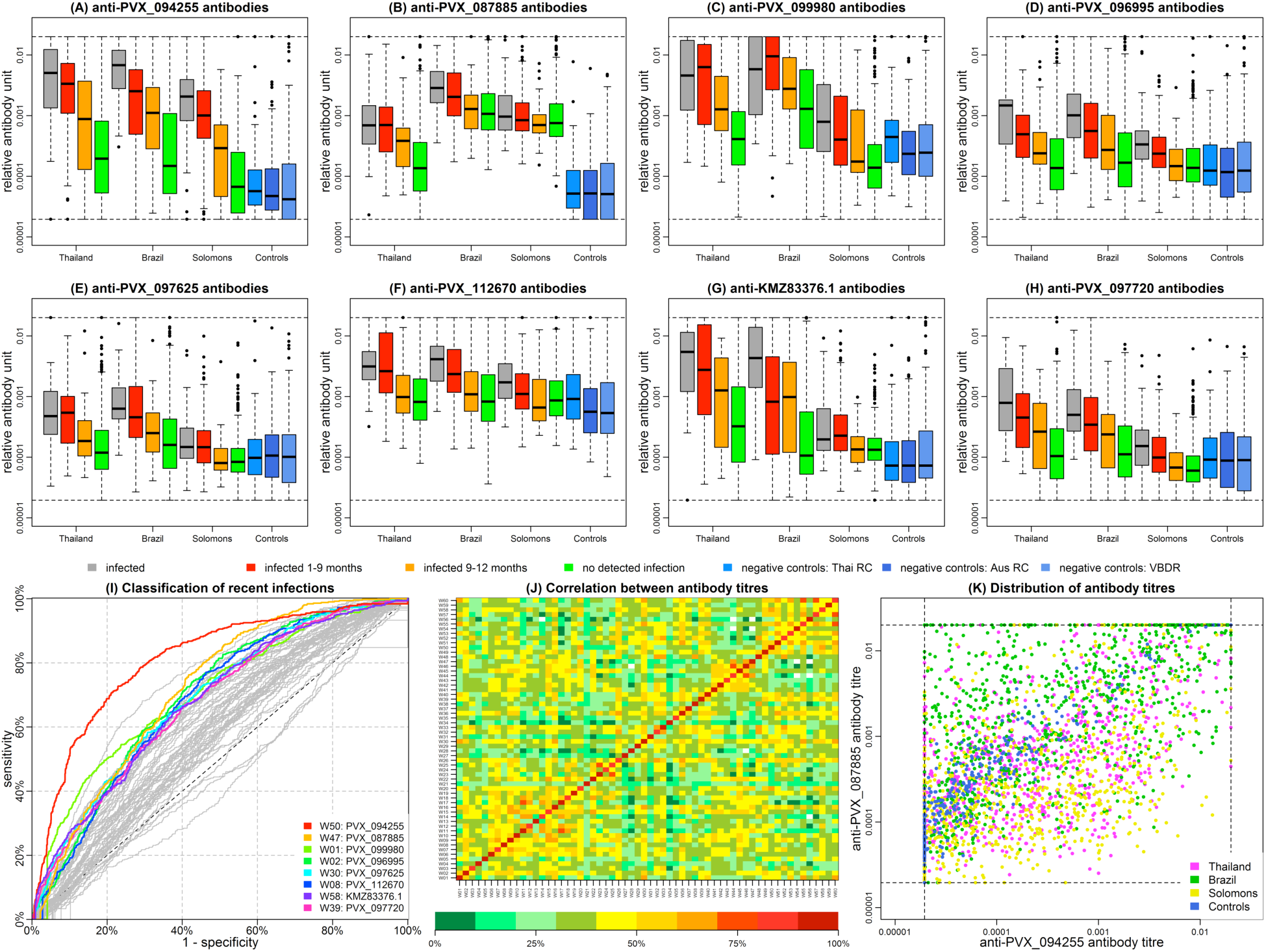
Validation phase. (A-H) Measured antibody response to 8 antigens on the Luminex® platform, stratified by geographical location and time since last PCR detected infection. (I) Receiver operating characteristic (ROC) curve for classifying individuals as infected within the last 9 months using a threshold antibody titre to a single antigen. Coloured curves correspond to those proteins in panels a-h. (J) Spearman correlation between antibody titres. Correlation coefficients are indicating by the colour scale, with red being 100% correlation and dark green 0% correlation. (K) 2-dimensional distribution of anti-PVX_099980 and anti-PVX_094255 antibody titres.

Recent infections were defined as PCR-confirmed infections occurring within the past 9 months. Using such a timeframe allows detection of likely hypnozoite carriers based on known relapse patterns (*20*), and enables analysis of individuals infected both before and after this period with the available data from the one-year long cohorts. Measurements of several individual proteins were able to classify individuals as infected with *P. vivax* within the past 9 months or not, with differing degrees of accuracy (Fig. 4i). Protein PVX_094255 (reticulocyte binding protein 2b, RBP2b) reached 74% sensitivity and 74% specificity when used alone. IgG levels to the 60 candidate SEMs were correlated (Fig. 4j) with different distributions evident between the three geographic regions (Fig. 4k). A linear discriminant analysis (LDA) classification algorithm was used to identify combinations of the 60 candidate SEMs that could accurately predict recent infections. A degree of redundancy was found with multiple combinations of *P. vivax* proteins able to accurately predict recent infection when included in panels of up to eight proteins (Fig.S6). The top eight most frequently identified proteins when used in combination were: PVX_094255 (RBP2b (*30*)), PVX_087885 (rhoptry associated membrane antigen, RAMA (*28*), putative), PVX_099980 (merozoite surface protein 1, MSP1 (*31*)), PVX_096995 (tryptophan-rich antigen (Pv-fam-a), PvTRAg_2 (*32*)), PVX_097625 (MSP8 (*33*), putative), PVX_112670 (unspecified product), KMZ83376.1 (erythrocyte binding protein II, EBPII (*34*)), and PVX_097720 (MSP3.10 (*35*)) (see Fig. 4i for ROC curves). PVX_112670 has previously been annotated as a tryptophan-rich antigen (PvTRAg_28 (*32*)). Table 2 and Table S2 show the individual ranking of all 60 proteins by AUC, note that it is not necessarily the proteins with the highest individual AUC that work best in combination.

As shown in Table 2, for the top protein PVX_094255 (RBP2b), there were two protein constructs, mapping to different regions of the protein (RBP2b161-1454 and RBP2b1986-2653), produced in two separate laboratories using different expression and purification methods (*30, 36*). Antibody levels to the two constructs of PVX_094255 (RBP2b161-1454 and RBP2b1986-2653) were highly correlated (spearman r=0·72, p<0·0001, all cohorts combined) and for this reason the construct that provided lower levels of classification accuracy (region encompassing aa1986-2653) was excluded from the top eight in order to increase the information content provided. RBP2b161-1454 induced very low levels of antibody reactivity in the malaria naïve control panels. When levels of background reactivity detected in the malaria naïve negative control panels were investigated in the other candidate SEMs, a significant negative association was observed between the mean antibody levels detected in the malaria naïve negative control panels (n=274 individuals, see Methods) and the ranking of the 60 SEMs determined from their individual receiver operating characteristic (ROC) curves (spearman r=0.6, p<0.0001, Fig. S10).

### Classification performance of an eight-protein SEM panel

Fig. 5a-d presents ROC curves for assessing the classification performance of the top panel of eight SEMs for identifying individuals with exposure to *P. vivax* within the prior 9 months. There are diminishing returns in classification performance as the number of proteins is increased, with a plateau of 80% sensitivity and 80% specificity reached in all three geographic regions with five proteins. The performance of the classification algorithm across the different sample types was assessed. The algorithm correctly classifies more than 98% of malaria naïve negative controls, but classification performance was poor for individuals who had their last blood-stage *P. vivax* infection between 9-12 months ago, with 30-60% of samples mis-classified (Fig. 6). Incorporating an individual’s age into the classification algorithm did not result in substantial increases in classification accuracy (Fig. S11). There was a significant relationship between background levels of antibody in malaria naïve controls and selection of top proteins in the validation phase (Fig. S10). However, removing data from these malaria naïve control participants did not cause any substantial reductions in classification accuracy for the top eight proteins (Fig. S12). The cohort studies were conducted in regions co-endemic for *P. vivax* and *P. falciparum*, although the total number of individuals experiencing a *P. falciparum* infection during the study period was low (n=19, 31, 22 for Thailand, Brazil and the Solomon Islands, respectively). No associations were observed between recent *P. falciparum* infections and antibody level to the top eight proteins (Fig. S13).

**Fig. 6.**
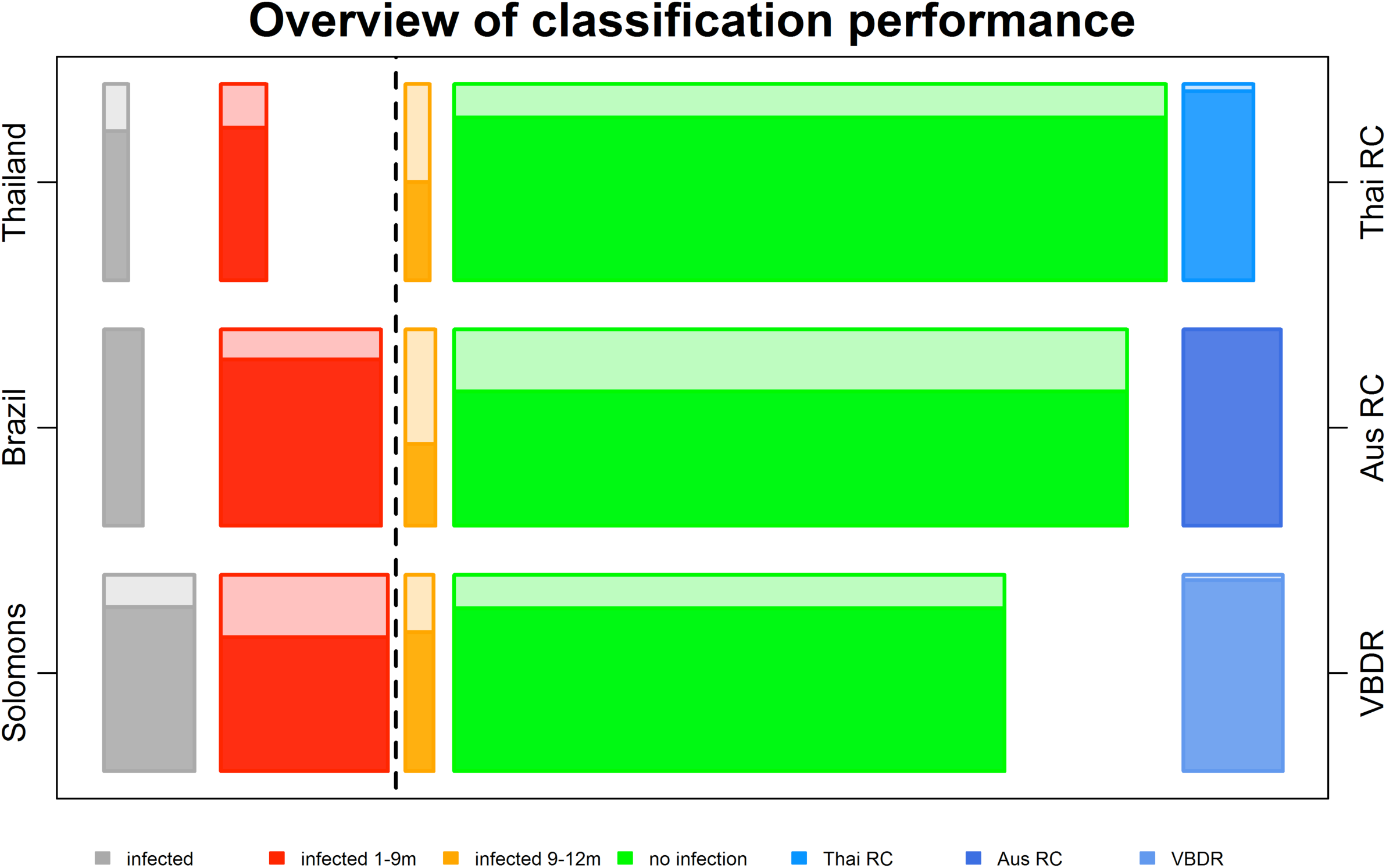
Breakdown of classification performance of the composite algorithm with target sensitivity and specificity of 80%. The size of each rectangle is proportional to the number of samples in each category (See Table 1). The coloured area represents the proportion correctly classified, and the shaded area represents the proportion mis-classified. Thai RC = Thai Red Cross, Aus RC = Australian Red Cross, VBDR = Volunteer Blood Donor Registry.

**Fig. 5.**
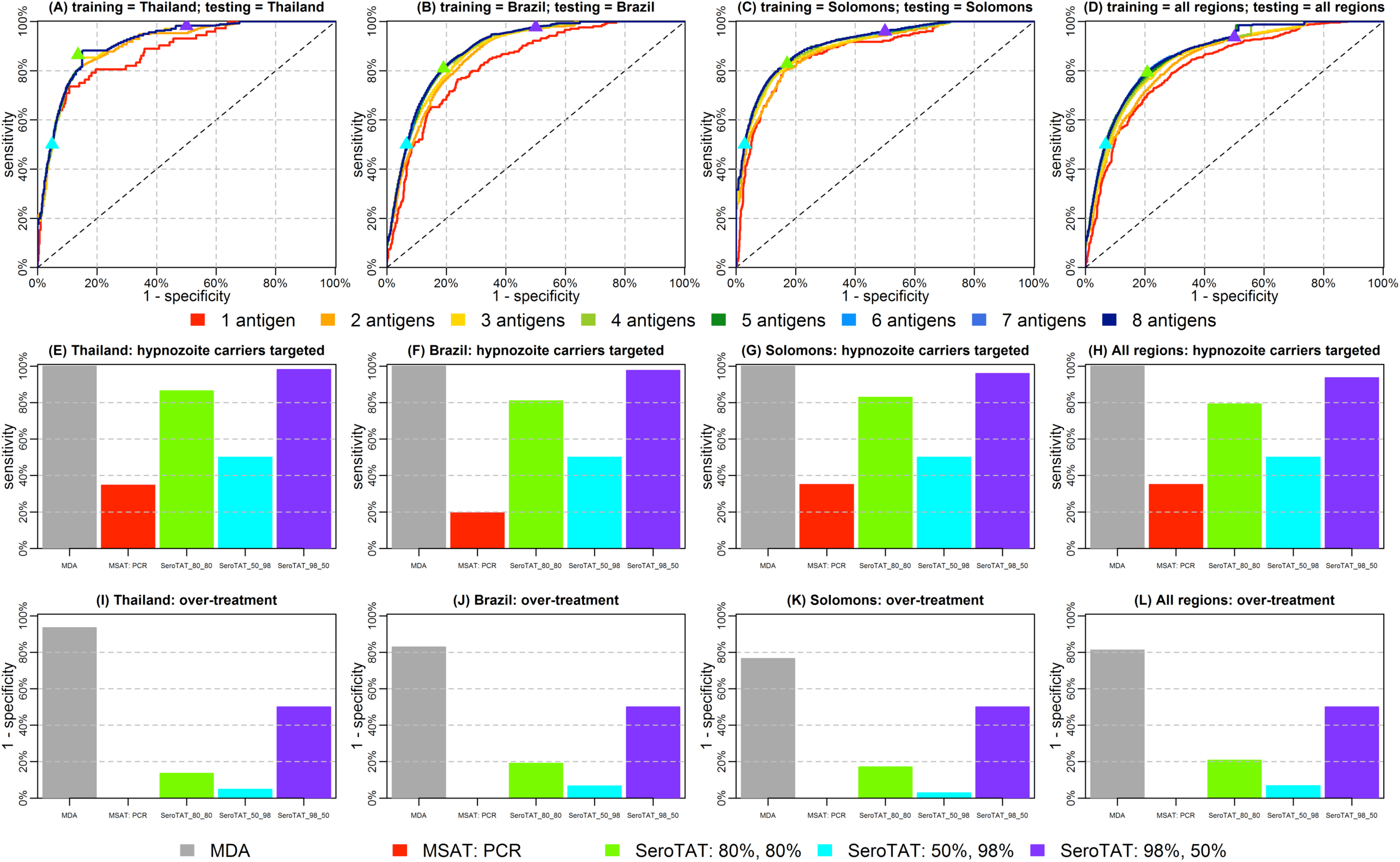
Performance of the top 8 *P. vivax* proteins for classifying recent infection and modelled ability to identify hypnozoites carriers. Cross-validated ROC curves generated from the composite algorithm classifier for identifying individuals with PCR detected infection in the last 9 months in (A) Thailand; (B) Brazil; (C) The Solomon Islands; and (D) all regions combined. The coloured triangles denote three different sensitivity and specificity targets for serological screening and treatment (SeroTAT) strategies. (E-H) Number of individuals targeted for primaquine treatment under a range of mass treatment strategies. (I-L) Proportion of individuals who had no exposure to *P. vivax* during the previous 9 months who would have been administered primaquine.

The performance of targeted treatment conducted with the top eight *P. vivax* SEMs using a seroTAT approach compared to both MDA and MSAT with PCR was modelled (Fig. 5). Both sensitivity and specificity were assessed. In an MDA campaign, all hypnozoite carriers are targeted (except those deemed ineligible due to G6PD deficiency), but > 80% of the population receive unnecessary primaquine. In contrast for MSAT with PCR, no individual was over-treated (it is assumed that an individual with blood-stage *P. vivax* is a likely hypnozoite carrier). However, only 20%-40% of likely hypnozoite carriers are targeted. These approaches were compared to seroTAT with the panel of top eight SEMs at 80% sensitivity and specificity (Fig. 5). Across the three cohorts (Thailand, Brazil, Solomon Islands), at least 80% of individuals with hypnozoites are targeted (outperforming MSAT with PCR), with 20% or less of the population treated unnecessarily (substantially less than an MDA approach). Given the balance between sensitivity and specificity, two alternate targets were also modelled: 50% sensitivity and 98% specificity, and 98% sensitivity and 50% specificity. The high specificity approach results in no overtreatment but only slightly outperforms MSAT in terms of identifying hypnozoite carriers (45% missed), and is likely not feasible for seroTAT. The high sensitivity approach identifies nearly all likely hypnozoites carriers and reduces overtreatment to approximately 50%, compared to >80% with MDA.

## Discussion

There is a clear and urgent need for new tools and strategies if we are to achieve malaria elimination in the Asia-Pacific by the year 2030 (*37*). *P. vivax*, the predominant *Plasmodium* species in the Asia-Pacific, presents a unique challenge to elimination due to the presence of undetectable hypnozoites, representing a hidden burden of infection that contributes to maintaining residual transmission. A novel panel of candidate serological exposure markers (“SEMs”) were identified and validated that allow for the detection of recent exposure to *P. vivax* within the prior 9-month period. Relapses from *P. vivax* infections are expected to occur at a frequency of every 1-2 months, with hypnozoites remaining in the liver for up to 1-2 years (*21, 22*). Although a few relapses may occur more than 1 year after the last blood-stage infection, the initial dormancy period for all *P. vivax* strains is less than 9 months with subsequent relapses occurring generally at shorter intervals (*20-22*). Apart from temperate ‘hibernans’ strains that are now restricted to the Korean Peninsula and have no primary infections, virtually all individuals who carry hypnozoites will thus have had a *P. vivax* blood-stage infection within the prior 9 months. To our knowledge, these SEMs represent the foundation for the first test that can indirectly identify likely hypnozoites carriers that could be targeted for treatment with liver-stage drugs.

We undertook an unbiased approach to choosing the best markers by starting the screen with a large panel of 342 *P. vivax* proteins, and strategically selecting those that can predict recent infection history based on immunogenicity and antibody half-lives. The final panel, which has a sensitivity and specificity of 80% at identifying individuals with PCR-detectable blood-stage infection in the last 9 months in three geographically distinct regions, incorporates antibody responses to eight *P. vivax* proteins: PVX_094255 (RBP2b161-1454), PVX_087885 (RAMA, putative), PVX_099980 (MSP1), PVX_096995 (tryptophan-rich antigen (Pv-fam-a)), PVX_097625 (MSP8, putative), PVX_112670 (unspecified product), KMZ83376.1 (EBPII), and PVX_097720 (MSP3.10). Most of these *P. vivax* proteins are not well described in the literature and only MSP1 and RAMA have previously been used, or suggested as, markers of exposure (*28, 38-40*). This highlights the success of our strategy in identifying novel exposure markers.

Individually, PVX_094255 (RBP2b161-1454) was able to classify exposure within the past 9 months with 74% sensitivity and 74% specificity. This may be sufficient for sero-surveillance and community-level targeting of preventative measures such as intensified vector control or focal test and treat (*19*). In such a scenario, lower sensitivity could be accounted for by increasing the sample size surveyed. Greater classification accuracy will be required for individual-level targeting for treatment with liver-stage drugs, for example in an elimination campaign using “serological testing and treatment” (seroTAT), where the goal is to treat all individuals who have hypnozoites or blood-stage *P. vivax* parasites. Combining PVX_094255 (RBP2b161-1454) with additional proteins resulted in improved classification performance, albeit with diminishing returns. The final panel of eight proteins obtained 80% sensitivity and 80% specificity across the three geographic regions tested.

Anti-malarial antibody responses are affected by a number of factors, most notably time since last infection, intensity of infection, and age. Antibody responses may also be associated with other factors such as genetic variation in the human host or parasite that are challenging to account for. When considering the relationship between time since last infection and measured antibody responses, key properties of the data are the high level of variation (some individuals with blood-stage infection had very low responses, and some negative controls had very high responses), and the high degree of correlation. The analytic pipeline was designed to account for this type of data, and the possibility that recent infections may best be identified by combinations of antibody responses. However, rather than past infections being identified by complex antibody signatures and sophisticated algorithms, a number of simple factors contributed to good classification performance: (i) *P. vivax* proteins that can identify recent infections when used individually also do well in combinations; (ii) there are diminishing returns in classification performance as new *P. vivax* proteins are added to combinations; (iii) there are diminishing returns to algorithmic complexity: simple algorithms such as logistic regression capture most of the signal in the data, with more complex classifiers such as random forests providing incremental improvements in performance; (iv) there is no single best combination of antigens, rather many antigens are interchangeable; (v) there was no substantial advantage to including an individual’s age once three or more antigens were included; and (vi) algorithms and combinations of antigens that provided good classification in one region also performed well in another region.

Although the performance of the pilot marker panel may seem low, several factors need to be taken into account when evaluating the potential performance of the SEMs. Firstly, given that individual episodes of *P. vivax* blood-stage can be relatively short (i.e. <4-6 weeks) (*41*), it is highly likely that some blood-stage infections may have been missed by qPCR in the cohorts used for validation of the SEMs as they had monthly active follow-up. In the analyses, such cases would be classified as false positive by the SEMs, when in reality the SEMs would have accurately detected these infections. Therefore, the real specificity of the test may well be higher. Similarly, a number of people (5-20%) with PCR-positive infections at the time of antibody measurement were missed. This may represent individuals with very recent infections who have not yet generated a significant rise in antibody titres (*42*). These missed, concurrent infections could however be detected using an ultra-sensitive antigen capture test. A combination of antibody detection and antigen-capture assays – implemented either on a single diagnostic platform or two tests run in parallel – would therefore further increase the sensitivity of the diagnostic approach. Lastly, the SEMs are designed for application in low-transmission malaria-endemic regions progressing towards elimination. Our data has previously shown that antibody responses are longer lasting in areas with higher transmission (*24*). In elimination contexts, overall (population) levels of immune responses to malarial antigens are low (*24, 43*), potentially increasing the difference between recently exposed and non-recently exposed individuals.

Although we anticipate optimal classification performance in near-elimination settings, our validation studies had to be conducted in low to moderate transmission settings, where the required sample sizes are operationally feasible. Indeed, in the three settings tested, classification performance was greatest in Thailand, the region with lowest transmission. The association of antibody levels to the 60 validation proteins with current *P. vivax* infections was less clear in the Solomon Islands compared to Thailand and Brazil, and the magnitude of the antibody response to these proteins was also generally lower. In contrast to the Thai and Brazilian cohorts that included individuals of all ages, the Solomon Islands study was a paediatric cohort. Antibody levels tend to increase with age in malaria-endemic regions (*13, 29*), and so the lower responses in the Solomon Islands may be due to the design of the cohort, which includes only children. However, the difference in antibody responses measured could also be influenced by other factors such as genetic diversity of *P. vivax* parasite proteins. Importantly, despite the lower magnitude antibody responses, the top eight SEMs could still accurately classify recent infections in the Solomon Islands.

To further improve classification performance using the top *P. vivax* proteins identified, modifications of the protein constructs resulting in reduced background in malaria naïve control samples would be advantageous. The data demonstrated that proteins with lower levels of antibody reactivity in these controls performed better at classifying recent exposure. Addition of further purification steps, testing of other protein expression systems, design of smaller protein fragments (for those currently full-length), investigations of antigenic diversity and strain-specificity of antibody responses, could all be considered. Further information could potentially be gained by looking at antibody subclasses and/or IgM in addition to total IgG responses. Whilst antibody subclass responses will likely correlate with total IgG, at least for the dominant subclass (*44*), the relationship with IgM is expected to be weaker (*44*) and acquisition and breadth of responses to malaria proteins has been shown to differ for IgG and IgM (*45*). Such optimisation and improvement of our assay may allow selection of a smaller panel of three to five proteins that will be able to reach the classification accuracy of our current panel of eight proteins. This would reduce the cost of running our SEMs in their current format (Luminex® assay) and also make a simpler point-of-care test feasible.

The SEMs current level of performance and their application in potential public health interventions such as seroTAT also needs to be seen in the context of the alternatives. MDA is a blanket approach, with liver-stage drugs given to all individuals within a defined region. Whilst this will effectively target all hypnozoite carriers and is predicted to achieve a high impact (but only if it includes an anti-hypnozoite drug (*5*)), it results in substantial overtreatment. In low transmission settings, up to 90% of people treated do not carry hypnozoites (i.e. the specificity of MDA can be as low at 6.5-23.3%, Fig. 5) and do not derive any direct benefit from a treatment that can have negative consequences in G6PD-deficient individuals. Therefore, few *P. vivax* endemic countries are considering MDA with primaquine. MSAT interventions on the other hand will miss most hypnozoite carriers and when implemented with light microscopy or RDT diagnosis will not achieve any impact on *P. vivax* transmission (*11*). Even with a highly sensitive molecular test, MSAT is likely to miss 64.9-80.5% of the people carrying hypnozoites (Fig. 5) with relapsing infections thus predicted to rapidly erode its effect (*5*). Testing and treatment with SEMs thus result in less overtreatment compared to MDA, whilst still targeting a higher proportion of likely hypnozoites carriers compared to MSAT by PCR.

Similar efforts to develop SEMs are currently underway for *P. falciparum* (*15*). Whilst *P. falciparum* does not have a latent liver-stage leading to relapse, detecting recent exposure may still provide a useful tool for prioritising the limited resources available to areas with residual transmission. As numerous countries are endemic for both *P. falciparum* and *P. vivax* (*1*), efforts to develop SEMs for both species should be coordinated where possible. In addition, there is some evidence of antibody cross-reactivity between *P. falciparum* and *P. vivax*: cross-reactivity for *P. falciparum* and *P. vivax* MSP5 was demonstrated by a competitive antibody assay in PNG (co-endemic for *P. falciparum* and *P. vivax*) (*46*). Five of the top eight proteins have *P. falciparum* orthologs (with protein sequence identity ranging from 20.7-37.4%) and thus need to be assessed for the impact of cross-reactivity in co-endemic regions. This is particularly important given that antibody longevity to the *P. falciparum* orthologs is unknown. No association was detected between recent *P. falciparum* infections and antibody levels to the SEMs, however the prevalence of *P. falciparum* infections in the validation cohorts was very low (0.3–0.6%). Given the relatively low sequence homology between *P. falciparum* and *P. vivax*, extensive cross-reactivity is unlikely. This is however not necessarily the case for *P. ovale, P. malariae* or *P. knowlesi* which are more closely related to *P. vivax* (protein sequence identity for top eight antigens 13.7-83.4%). On a programmatic level, limited cross-reactivity between *P. vivax, P. falciparum, P. ovale* or *P. malariae* may not be an issue in areas outside Africa. It has been well documented that rates of *P. vivax* relapse after *P. falciparum* episodes or infections are very high (*47, 48*), sometimes as high as following a *P. vivax* infection. This indicates that the presence of a *P. falciparum* infection is a marker of *P. vivax* hypnozoite carriage and it has therefore been suggested that in co-endemic areas all *P. falciparum* patients should receive anti-*P. vivax* relapse therapy (*47*). The same is likely to apply to *P. malariae* and *P. ovale* infections, which regularly occur in mixed species infections (*49*), and of which the latter also has dormant hypnozoites. Given the marked difference in patterns of exposure to the zoonotic *P. knowlesi* (*50*), cross-reactivity between *P. vivax* and *P. knowlesi* would be significantly more problematic (*51*).

In summary, the novel SEMs for *P. vivax* can accurately predict recent infection within the last 9 months. As most hypnozoites in tropical regions are expected to relapse within 9 months of a previous blood-stage infections (*52*), these markers can indirectly identify the individuals with the highest risk of harbouring hypnozoites. Past efforts for developing serology as a measure of exposure to *P. vivax* have only detected historical changes over time (*53*); we have shown that a carefully selected panel of SEMs can specifically detect recent exposure and could be used in a programmatic setting (“surveillance as an intervention” (*54*)). Application of our SEMs for seroTAT holds promise for an effective elimination strategy using primaquine or tafenoquine to target dormant hypnozoites. Given the risk of haemolysis in G6PD-deficient individuals treated with primaquine or tafenoquine, our tool has the potential to ensure such elimination strategies are targeted and therefore a safer, more acceptable and more effective option in malaria-endemic communities.

## Materials and Methods

### Study design

The goal of the study was to identify then validate suitable candidate *P. vivax* proteins that could be used as serological markers of recent exposure to *P. vivax* infections. This study was conducted in two parts: i) an antigen discovery phase utilising samples from two longitudinal cohorts that followed symptomatic *P. vivax* malaria patients over 9 months, and ii) a validation phase utilising samples from three one-year long observational cohort studies. The sample size was therefore predefined by availability of suitable plasma samples with matching epidemiological and molecular data.

### Study populations: antigen discovery phase

Patients with confirmed *P. vivax* malaria were enrolled from local malaria clinics and hospitals in two sites in 2014: Tha Song Yang, Tak Province, Thailand, and Manaus, Amazonas State, Brazil. After receiving anti-malarial drug treatment according to respective National Treatment Guidelines, and providing written informed consent and/or assent, blood samples were taken over a period of 9 months as previously described (*24*) (Fig. 1a, Table 1). Presence or absence of *Plasmodium spp.* parasites during follow-up was determined by both microscopy and quantitative PCR. Blood samples were collected at enrolment and week 1, then every 2 weeks for 6 months, then every month until the end of the study. A subset of enrolled participants who had no evidence of recurrent parasitaemia during follow-up was selected for antibody measurements (n=32 Thai patients, n=33 Brazilian patients).

The Ethics Committee of the Faculty of Tropical Medicine, Mahidol University, Thailand, approved the Thai study (MUTM 2014-025-01 and 02). The Ethics Review Board of the Fundação de Medicina Tropical Dr. Heitor Vieira Dourado (FMT-HVD) (957.875/2014) approved the Brazilian antigen discovery study. The Human Research Ethics Committee (HREC) at the Walter and Eliza Hall Institute of Medical Research (WEHI) approved samples for use in Melbourne (#14/02).

### Study populations: validation phase

Yearlong observational cohort studies were conducted over 2013-2014 in three sites: Kanchanaburi/Ratchaburi provinces, Thailand (*14*), Manaus, Brazil (Monteiro et al., in preparation), and the island of Ngella, Solomon Islands (Quah et al., in preparation) (Fig. 1b, Table 1). A total of 999 volunteers were enrolled from Thailand and sampled every month over the yearlong cohort, with 14 active case detection visits performed in total. 829 volunteers attended the final visit. A total of 1,274 residents of all age groups were enrolled from Brazil and sampled every month over the year-long period, with 13 active case detection visits performed in total. 928 volunteers attended the final visit with plasma from 925 available. 1,111 children were enrolled from the Solomon Islands, with 860 used for final analysis of the cohort (after exclusion of children who withdrew, had inconsistent attendance or failed to meet other inclusion criteria). The children were sampled monthly, with 11 active case detection visits in total. Of the 860 children, 754 attended the final visit. For all three studies, at each visit volunteers completed a brief survey outlining their health over the past month (to determine the possibility of missed malarial infections), in addition to travel history and bed net usage. A finger-prick blood sample was also taken and axillary temperature recorded. Passive case detection was performed throughout the yearlong period.

The Ethics Committee of the Faculty of Tropical Medicine, Mahidol University, Thailand approved the Thai cohort (MUTM 2013-027-01). The Brazilian study was approved by the FMT-HVD (51536/2012), by the Brazilian National Committee of Ethics (CONEP) (349.211/2013) and by the Ethics Committee of the Hospital Clínic, Barcelona, Spain (2012/7306). The National Health Research and Ethics Committee of the Solomon Islands Ministry of Health and Medical Services (HRC12/022) approved the Solomon Islands study. The HREC at WEHI approved samples for use in Melbourne (#14/02).

### Study populations: malaria naïve control panels

Three panels of malaria naïve control plasma samples were collected from individuals with no known recent exposure to malaria (Table 1). Samples were as follows: 102 individuals from the Volunteer Blood Donor Registry (VBDR) in Melbourne, Australia, 100 individuals from the Australian Red Cross (ARC), Melbourne, Australia, and 72 individuals from the Thai Red Cross (TRC), Bangkok, Thailand. Standard TRC screening procedures exclude individuals from donating blood if they have had a confirmed malaria infection within the previous 3 years or have travelled to malaria-endemic regions within the past year. The HREC at WEHI approved collection and/or use of these control samples (#14/02).

### Procedures

Blood samples were collected by finger prick into EDTA tubes. Blood was separated into packed red cells and plasma at the field site. Packed red cells were stored at −20°C and plasma at −80°C prior to use in molecular and serological assays, respectively. Malaria parasites were detected by qPCR as previously described (*55, 56*).

### Antibody measurements

*P. vivax* malaria proteins were expressed using the wheat germ cell-free (WGCF) system (CellFree Sciences, Yokohama, Japan) at either Ehime University or CellFree Sciences. For the antigen discovery phase, 307 *P. vivax* protein constructs were expressed at a small-scale with a biotin tag and antibodies measured using the AlphaScreen™ assay, as previously described (*24*). Plasma samples from weeks zero (enrolment), 12, 24 and 36 in the subset of volunteers described above were used (n=32 Thai patients, n=33 Brazilian patients). The raw AlphaScreen™ data was converted to relative antibody units based on plate-specific standard curves, with final units ranging from 0-400 (seropositivity was defined as a relative antibody unit greater than 0).

For the validation phase, down-selected proteins were produced at a high yield using dialysis-based refeeding WGCF methods and purified on an affinity matrix using a His-tag. The purified proteins were stored and shipped in the following buffer: 20 mM Na-phosphate pH 7·5, 0·3 M NaCl, 500 mM imidazole and 10% (v/v) glycerol. Protein yields and purity were determined using SDS-PAGE followed by Coomassie Brilliant Blue staining using standard laboratory methods. An additional 20 *P. vivax* proteins known to be highly immunogenic were also included at the validation phase and were produced as previously described (*26-28, 30, 57, 58*) or using previously described methods (*59*) (see Table 2 and Table S2). The *P. vivax* proteins were then coupled to COOH microspheres as previously described (*13*). Protein-specific IgG antibody levels were measured in a multiplexed Luminex® assay as previously described (*29*). Plasma samples from the last visit of the three year-long observational cohort studies were used. Median fluorescent intensity (MFI) values from the Luminex®-200 were converted to arbitrary relative antibody units based on a standard curve generated with a positive control plasma pool from highly immune adults from Papua New Guinea (PNG) (*29*).

### Statistical modelling – antigen discovery phase

The change in measured antibody responses following *P. vivax* infection in patients in the antigen discovery phase was analysed using mixed-effects linear regression models, as previously described (*24*). Estimated half-lives for 307 *P. vivax* proteins were previously reported (*24*), with an additional 35 proteins included for this study. All estimated half-lives are provided in Supplementary Table S1. Antibody responses were measured at 0, 3, 6 and 9 months using the AlphaScreen™ assay, and the model was only fitted to individuals who were seropositive at baseline. Denote *Aijk* to be the antibody titre in participant *i* to protein *j* at time *tk* which can be described by the following linear model:

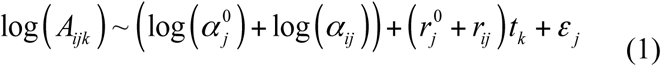

where *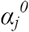* is the geometric mean titre (GMT) at the time of infection; log(*αij)* is a random effect accounting for the difference between participant *i*’s initial antibody titre and the population-level GMT; *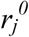* is the average rate of decay of antibodies to protein *j* in the population; *r*_*ij*_ is a random effect for the difference between the decay rate of individual *i* with the population-level average; and *ε*_*j*_ ∼ N(0, *σ*_*m,j*_) is a Normally distributed error term. In particular it is assumed that the random effects for the initial antibody titres are Normally distributed: log(*α*_*ij*_) ∼ N(0, *σ*_*A,j*_), and that the random effects for the variation in decay rates are also Normally distributed: *r*_*ij*_ ∼ N(0, *σ*_*r,j*_).

The linear regression model for the decay of antibody titres described in equation (S1) has three sources of variation: (i) variation in initial antibody response following infection; (ii) between individual variation in antibody decay rate; and (iii) measurement error. Notably, all these sources of variations are described by Normal distributions so their combined variation will also be described by a Normal distribution. Therefore, *x*_*ij*_ = log(*A*_*ij*_), the expected log antibody titre in individual *i* to protein *j* at time *t,* can be described by the following distribution:

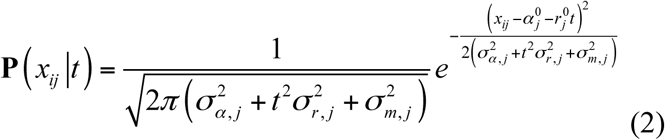

The probability distribution for the time since infection *t* given measured antibody titre *xij* can be calculated by inverting equation (1) using Bayes rule (*60*), allowing estimation of the probability that the time since last infection was less than 9 months.

It was possible to select >10^42^ subsets from the panel of 142 prioritised SEMs in the antigen discovery phase. A simulated annealing algorithm was therefore used to explore this high-dimensional space and select optimal combinations of proteins (*26, 61*). Combinations of proteins were chosen to optimise the likelihood that sampled antibody responses were correctly classified as having occurred within 9 months of infection.

### Statistical modelling – validation phase

Individuals in the validation phase were classified into two categories depending on whether they had PCR-detectable blood-stage *P. vivax* infections within the 9 months prior to measurement of antibody responses (Table 1). In theory it is possible to provide quantitative estimates of the time since last infection, however a more useful outcome in practice is whether an individual has been infected within some past-time period. 9 months was selected due to the biology of *P. vivax* hypnozoite relapses as described in the Introduction (highest incidence of relapse within the first 9 months following mosquito bite infection (*20*)), and because this threshold fell within a time period for which data were available (the three longitudinal cohorts had follow-up for 12 months).

Initial classification performance of antibody responses to the 60 down-selected SEMs to classify recent *P. vivax* infections was assessed using a linear discriminant analysis (LDA). There were 1,770 ways of choosing two out of 60 proteins, 34,220 ways of choosing three proteins, 487,635 ways of choosing four proteins, and 5,461,512 ways of choosing five proteins. All combinations up to size four were exhaustively evaluated to optimise classification performance for the three sensitivity and specificity targets described in the Results. To investigate combinations of size five, the 100 best combinations of four proteins for each of the three TPPs were identified. All possible remaining proteins were added to each of these 300 combinations, and the classification performance of all of these combinations of size five was assessed. A similar procedure was implemented to investigate classification performance of combinations of proteins up to size eight. Fig. S6 shows an overview of classification performance.

After determining the highest ranking SEMs using the LDA classifier, the top eight candidates were further assessed using a range of other classification algorithms, including logistic regression, quadratic discriminant analysis (QDA), decision trees, and random forests (*62*). Decision trees were implemented using the rpart R package. Random forests were implemented using the randomForest R package. Two novel classification algorithms incorporating a statistical model of antibody decay over time were also developed (see Supplementary Materials and Methods). These algorithms were combined into a composite algorithm so that the optimal algorithm and selection of proteins was selected for each target of sensitivity and specificity. All classification algorithms were cross-validated using 500 randomly sampled, disjoint training and testing subsets.

## Data Availability

All data is available within the figures and tables of the results and supplementary materials. Antibody half-lives and parameters used to generate Figure 3 are in Table S1.

## Code Availability

Code for statistical modeling can be provided upon request.

## Supporting information

## Supplementary Materials

### Materials and Methods

Fig. S1. Association between measured antibody titre and time since infection. Fig. S2. Dynamics of multiple antibodies.

Fig. S3. Estimates of (A) antibody half-life, and (B) the variance of each antibody response for 104 *P. vivax* proteins in the antigen discovery phase.

Fig. S4. Increasing the maximum number of proteins allowed in a panel leads to diminishing increases in likelihood and classification performance.

Fig. S5. Measured antibody responses to 60 proteins on the Luminex® platform, stratified by geographical location and time since last PCR detected infection.

Fig. S6. Performance of LDA classifier for combinations of proteins up to size eight for identifying individuals with blood-stage *P. vivax* infection within the last 9 months.

Fig S7. Network visualization of antigens.

Fig S8. Variable importance plot generated by a random forests algorithm for identify antigens that contribute to accurate classification.

Fig. S9. Receiver operating characteristic (ROC) curves depicting comparison of cross-validated classification performance for the seven classification algorithms considered.

Fig. S10. Receiver operating characteristic (ROC) curve for the composite classification algorithm.

Fig. S11. Association between background reactivity in non-malaria exposed controls and ranking of candidate SEMs.

Fig. S12. Assessment of incorporating information on an individual’s age into a Random Forests classifier.

Fig. S13. Assessment of the role of removing data from negative control participants on classification performance using a Random Forests classifier.

Fig. S14. Association between measurements of our top eight *P. vivax* proteins and time since last PCR detected blood-stage *P. falciparum* infection.

Table S1. Estimated antibody half-lives from antigen discovery phase.

Table S2. Purified *P. vivax* proteins used in the validation phase and their individual performance, complete list.

Table S3. Association of antibody level with current *P. vivax* infection.

## Acknowledgments

We gratefully acknowledge the extensive field teams that contributed to sample collection and qPCR assays, including Andrea Kuehn, Yi Wan Quah, Piyarat Sripoorote, and Andrea Waltmann. We thank all the individuals that participated in each of the studies, and thank the Australian and Thai Red Cross for donation of plasma samples. We thank the Volunteer Blood Donor Registry at WEHI for donation of plasma samples, and Lina Laskos and Jenni Harris for their collection and advice. We thank Christopher King (Case Western Reserve University) for provision of the PNG control plasma pool. Melanie Bahlo (Walter and Eliza Hall Institute) is thanked for her advice on algorithm development.

## Author contributions

RJL, MW, TT and IM designed the study. WN, WM, JK, ML, JS and IM conducted the cohort studies. ET, MM, MH, FM, W-HT, JH, CH, CEC and TT expressed the proteins. RJL, ET, MM, JB, and S-JL performed antibody measurements. RJL, CSNL-W-S and MW did the data management and analysis. MW performed modelling. RJL, MW and IM wrote the draft of the report. LJR, CP, DLD, XCD and IJG provided expert advice. All authors contributed to data interpretation and revision of the report.

