## Supplementary material for "Development and validation of serological markers for detecting recent exposure to *Plasmodium vivax* infection"

#### Supplementary Methods and Results

#### 1. Antigen discovery phase

##### 1.1. Linear model for decay of antibody response

The half-lives of the antibody responses to the 307 *P. vivax* proteins in the antigen discovery phase were estimated using previously described methods (Longley 2017). Estimated half-lives were calculated for an additional 35 proteins for this manuscript. In brief, 32 Thai individuals and 33 Brazilian individuals were followed longitudinally after a clinical episode of *P. vivax*. These individuals were treated with primaquine to prevent relapses, and the absence of any blood-stage *Plasmodium* parasites during follow-up was confirmed by testing blood samples by PCR. Antibody responses were measured at 0, 3, 6 and 9 months using the AlphaScreen assay. Denote $A_{ijk}$ to be the antibody titre in participant $i$ to protein $j$ at time $t_k$ which can be described by the following linear model:

$$\log\left(A_{ijk}\right) \sim \left(\log\left(\alpha_j^0\right) + \log\left(\alpha_{ij}\right)\right) + \left(r_j^0 + r_{ij}\right) t_k + \varepsilon_j \qquad \text{(S1)}$$

where $\alpha_j^0$ is the geometric mean titre (GMT) at the time of infection; log( $\alpha_{ij}$ ) is a random effect accounting for the difference between participant $i$'s initial antibody titre and the population-level GMT; $r_j^0$ is the average rate of decay of antibodies to protein $j$ in the population; $r_{ij}$ is a random effect for the difference between the decay rate of individual $i$ with the population-level average; and $\varepsilon_j \sim N(0, \sigma_{m,j})$ is a Normally distributed error term. In particular we assume that the random effects for the initial antibody titres are Normally distributed: $\log(\alpha_{ij}) \sim N(0, \sigma_{A,j})$, and that the random effects for the variation in decay rates are also Normally distributed: $r_{ij} \sim N(0, \sigma_{r,j})$.

The model was only fitted to individuals who were seropositive at baseline. This model also generated an estimate of the total variation in the data (arising from initial antibody level measured, the rate of antibody decay and the measurement error). Table S1 provides the estimated antibody half-lives and relevant parameters for all 342 proteins (see separate excel file).

##### 1.2. Estimation of time since infection using antibodies to a single *P. vivax* protein

The linear regression model for the decay of antibody titres described in equation (S1) has three sources of variation: (i) variation in initial antibody response following infection; (ii) between individual variation in antibody decay rate; and (iii) measurement error. Notably, all these sources of variations are described by Normal distributions (Figure S1a) so their combined variation will also be described by a Normal distribution. Therefore, $x_{ij} = \log(A_{ij})$, the expected log antibody titre in individual $i$ to protein $j$ at time $t$, can be described by the following distribution:

$$x_{ij} \sim N\left(\alpha_j^0 + r_j t, \sigma_{\alpha,j}^2 + t^2\sigma_{r,j}^2 + \sigma_{m.j}^2\right) \qquad \text{(S2)}$$

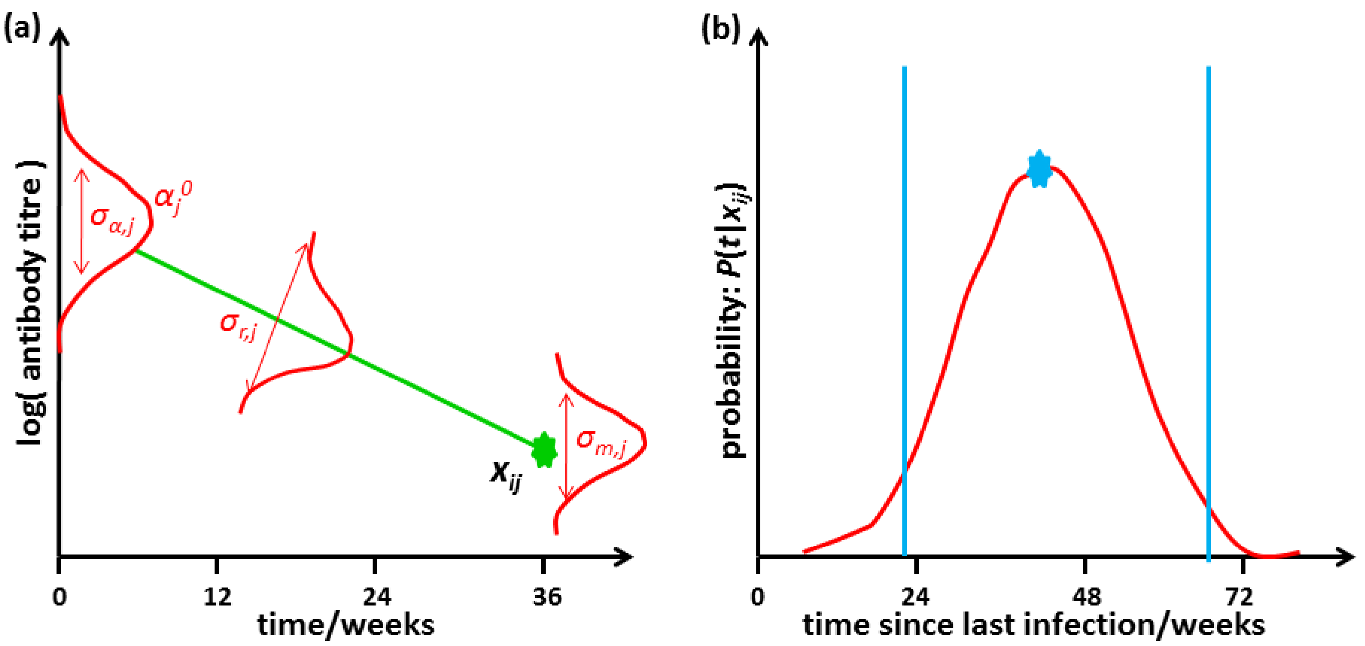


**Figure S1:** Association between measured antibody titre and time since infection. **(a)** There are three sources of variation in the log antibody titre $x_{ij}$ measured at time $t$ since last infection: (i) variation in initial antibody titre; (ii) between individual variation in antibody decay rate; and (iii) measurement error. **(b)** Given estimates of the sources of variation, we can estimate the distribution of the time since last infection. The maximum likelihood estimate and the 95% confidence intervals of our estimate are indicated in blue.

The probability distribution of the expected antibody titre in individual $i$ to protein $j$ at time $t$ is given by the following distribution:

$$\mathbf{P}\left(x_{ij}\middle|t\right)=\frac{1}{\sqrt{2\pi\left(\sigma_{\alpha,j}^2+t^2\sigma_{r,j}^2+\sigma_{m,j}^2\right)}}e^{-\frac{\left(x_{ij}-\alpha_j^0-r_j^0t\right)^2}{2\left(\sigma_{\alpha,j}^2+t^2\sigma_{r,j}^2+\sigma_{m,j}^2\right)}} \qquad \text{(S3)}$$

Note that we have $x_{ij}$ ϵ (-∞, +∞), as $x_{ij}$ denotes the log antibody titre and measurements of antibody titre are assumed to be positive. The probability distribution for the time since infection $t$ given measured antibody titre $x_{ij}$ can be calculated by inverting equation (S3) using Bayes rule (see also Borremans 2016).

$$\mathbf{P}\left(t\middle|x_{ij}\right)=\frac{\mathbf{P}\left(x_{ij}\middle|t\right)\mathbf{P}(t)}{\mathbf{P}\left(x_{ij}\right)} \qquad \text{(S4)}$$

The time since last infection will have a lower bound of zero. We can choose to impose an upper bound of either the individual's age $a$ or positive infinity. Choosing positive infinity allows us to better handle the case where an individual was never infected – the low measured antibody titres will be consistent with a very large time since last infection, possibly greater than the age of the individual. Therefore we should only restrict $t$ to the interval (0, $a$) if we know for certain that the individual has been infected. In practice, we choose some large time $t_{\max}$ = 100 years for our upper bound. We assume ***P***( $t$ ) denotes a uniform distribution on the interval (0, $t_{\max}$). ***P***( $x_{ij}$ ) is a normalising constant which is calculated via numerical integration to ensure that ***P***( $t$ | $x_{ij}$ ) denotes a probability distribution.

Equation (S4) provides a probability distribution for the time since last infection. For the purposes of a diagnostic test we may be more interested in obtaining a binary classification, e.g. was the individual infected within the last 9 months? We denote $z_i$ to be an indicator variable denoting whether individual $i$ was infected within the last 9 months ($z_i$ = 1) or not ($z_i$ = 0). The probabilities of these two events can be calculated as follows:

$$\begin{aligned}\mathbf{P}\left(z_i=1\middle|x_{ij}\right)&=\int_0^9\mathbf{P}\left(t\middle|x_{ij}\right)dt\\ \mathbf{P}\left(z_i=0\middle|x_{ij}\right)&=\int_9^{t_{\max}}\mathbf{P}\left(t\middle|x_{ij}\right)dt\end{aligned} \qquad \text{(S5)}$$

##### 1.3. Estimation of time since infection using antibodies to multiple *P. vivax* proteins

The methods above describe how measurements of antibody responses to a single protein can be used to estimate the time since last infection. However, in practice there is too much noise to make an accurate estimate of time since last infection with a single protein. Increasing the number of measured antibodies can increase the information content in our data allowing us to obtain more accurate estimates of time since last infection. In particular, selecting antibodies with a range of half-lives may increase our power to resolve infection times more accurately.

Figure S2 shows a schematic of the dynamics of antibodies to two proteins. We have rapidly decaying antibody 1 and slowly decaying antibody 2. At baseline, antibody titres are likely to be correlated, so we assume that initial titre following infection is described by a multivariate Normal distribution with covariance matrix $\Sigma_\alpha$. The between individual rates of antibody decay may also be correlated (i.e. all antibody titres may decay particularly quickly in some individuals) so we also assume that decay rates are described by a multivariate Normal distribution with covariance matrix $\Sigma_r$. Finally, there will be measurement error associated with each antibody. In particular, we assume the measurement errors between different antibodies are independent so that the total measurement error can be described by a multivariate Normal distribution with diagonal covariance matrix $\Sigma_m$. For the case with measurements of antibodies to $J$ proteins, we have:

$$\mathbf{P}(x_i|t) = (2\pi)^{-\frac{J}{2}} \left|\Sigma_\alpha + t^2\Sigma_r + \Sigma_m\right|^{-\frac{1}{2}} e^{-\frac{1}{2}\left(x_i - \alpha^0 - r^0 t\right)^T \left(\Sigma_\alpha + t^2\Sigma_r + \Sigma_m\right)^{-1}\left(x_i - \alpha^0 - r^0 t\right)} \tag{S6}$$

The time since last infection can be estimated given the multivariate probability distribution for the measured vector of antibody titres $x_i$ and the classification probability for the 9 month threshold can be estimated as follows:

$$\begin{aligned}\mathbf{P}(z_i = 1|x_i) &= \int_0^9 \mathbf{P}(t|x_i)\,dt \\ \mathbf{P}(z_i = 0|x_i) &= \int_9^{t_{\max}} \mathbf{P}(t|x_i)\,dt\end{aligned} \tag{S7}$$

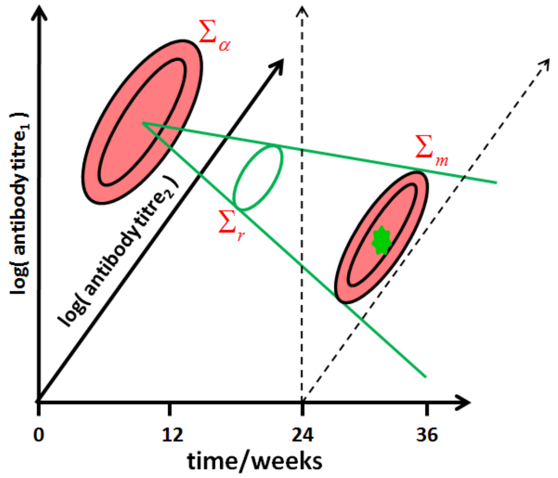


**Figure S2:** Dynamics of multiple antibodies. The variance in initial antibody titre, antibody decay rates and measurement error are now described by covariance matrices which account for the correlations between antibodies.

##### 1.4. Identifying optimal combinations of *P. vivax* proteins using simulated annealing

Machine learning algorithms take data from a large number of streams and identify which data streams have the most signal for classifying output. Such methods typically involve a greedy algorithm that will provide a good but not necessarily optimal solution. Greedy algorithms take the next best step, i.e. including the next protein that gives the biggest increase in predictive power. As such they may provide a locally optimal solution but not necessarily a globally optimal solution. Simulated annealing algorithms provide an alternative to greedy algorithms that usually provide a higher likelihood of obtaining a globally optimal solution (Kirkpatrick 1983).

Here we describe how a simulated annealing algorithm can be applied to the method for estimating time since infection described above, to select combinations of proteins that maximise the likelihood of correctly classifying infections within the last 9 months. Assume that $J$ measurements of antibodies are available. We want to select some subset of these that maximises predictive power. Denote $y$ to be a vector of 0's and 1's indicating whether the $j^{th}$ antibody is included in our panel. Thus for example with $J = 9$ we may have

$$y = (0,0,1,1,0,1,0,0,1) \tag{S8}$$

The vector of binary states depicted in equation (S8) will correspond to a vector of antibody measurements from individual *i* as follows:

$$x_i = \left( x_{i,3}, x_{i,4}, x_{i,6}, x_{i,9} \right) \qquad \text{(S9)}$$

Let $z_i$ be an indicator denoting whether individual *i* was infected in the last 9 months ( $z_i = 1$ ) or not ( $z_i = 0$ ). Given data from *I* individuals on measured antibody responses, we can calculate the probability that the individual was infected within the last 9 months $\boldsymbol{P}( z_i = 1 \mid x_i )$ or greater than 9 months ago $\boldsymbol{P}( z_i = 0 \mid x_i )$. We can then write down the likelihood of the data as follows:

$$L(y) = \prod_{i=1}^{I} \mathbf{P}\left(z_i = 1 \middle| x_i\right)^{z_i} \mathbf{P}\left(z_i = 0 \middle| x_i\right)^{1-z_i} \qquad \text{(S10)}$$

The challenge is to select a binary vector *y* corresponding to a combination of proteins that maximises the likelihood in equation (S10) and thus has the highest likelihood of correctly classifying infections according to whether they occurred in the last 9 months.

If we have *J* proteins, there are $2^J$ combinations of proteins. For $J > 15$ it is not computationally feasible to test all possible combinations. We therefore utilise a simulated annealing algorithm for exploring the state space of combinations and identifying the optimal combinations subject to various constraints (e.g. enforcing a maximum of $J_{max} = 10$ proteins to a panel).

Here we describe a simulated annealing algorithm with *M* iterations and an exponential cooling schedule such that the temperature at iteration $m \leq M$ is $T_m = e^{-\left(\frac{M-m}{M}\right)}$. It is also assumed that $J_{\max}$ is the maximum number of proteins that can be considered in a panel.

- At iteration *m* we have binary state vector $y_m$ with likelihood $L( y_m )$.
- Generate new binary state vector $y_{m+1}$ by switching two randomly selected binary states.
- Reject the new state if $\sum_{j=1}^{J} y_{j,m+1} > J_{\max}$
- Accept the new state with probability $\min\left( 1, e^{-\left(\frac{L(y_m) - L(y_{m+1})}{T_{m+1}}\right)} \right)$

The algorithm was repeated 100 times with different starting points and $M = 30000$ iterations. This resulted in 100 possibly different estimates of optimal combinations of proteins.

#### 1.5. Additional results

The output of the search of the space of protein combinations is summarised in Figure 3E of the main manuscript. Additional properties of the 104 *P. vivax* proteins considered in the second round of the antigen discovery phase are shown in Figure S3.

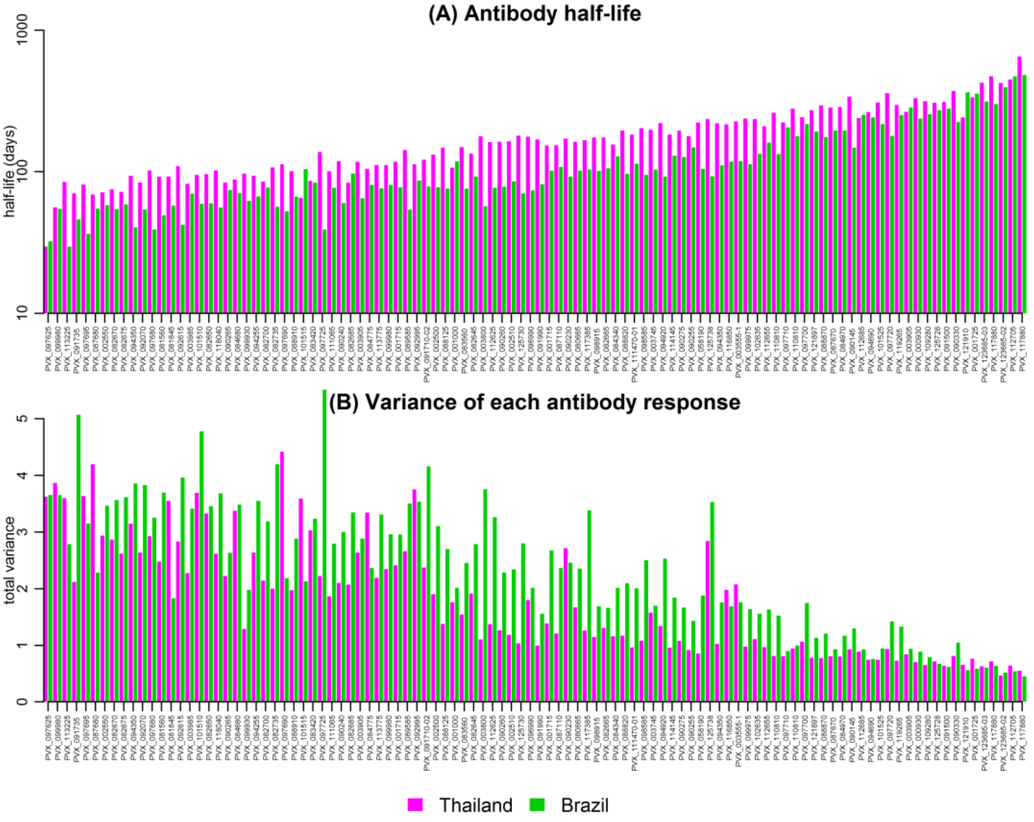


**Figure S3:** Estimates of (A) antibody half-life, and (B) the variance of each antibody response for 104 *P. vivax* proteins in the antigen discovery phase.

A property of the simulated annealing based search strategy applied to this data set is that, on average, the most predictive proteins are selected first, much like what would be expected for a greedy algorithm. This leads to a phenomenon of diminishing returns, where the addition of each new protein causes sequentially smaller improvements in likelihood and classification performance (Figure S4).

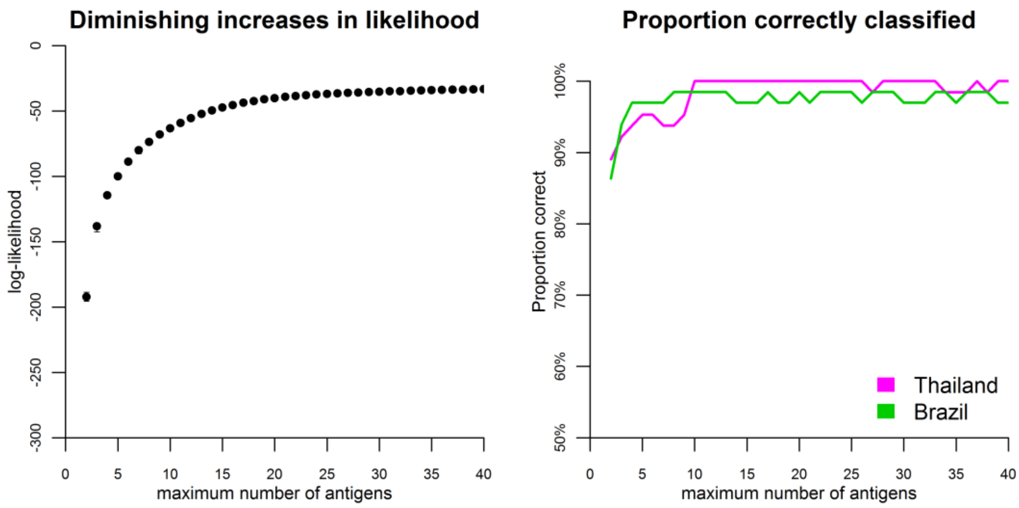


**Figure S4:** Increasing the maximum number of proteins allowed in a panel leads to diminishing increases in likelihood and classification performance. Note that classification performance was assessed using the same training and testing data set, i.e. without cross-validation.

### 2. Validation phase

#### 2.1. Supplementary data on all proteins

Table S2 shows the association of antibody level with current *P. vivax* infections (see separate excel file). Figure S5 shows the distribution of measured antibody responses to the 60 proteins used in the validation phase in the Thai, Brazilian, Solomon Islands and negative control cohorts, as well as the association with time since last PCR-detected blood-stage *P. vivax* infection.

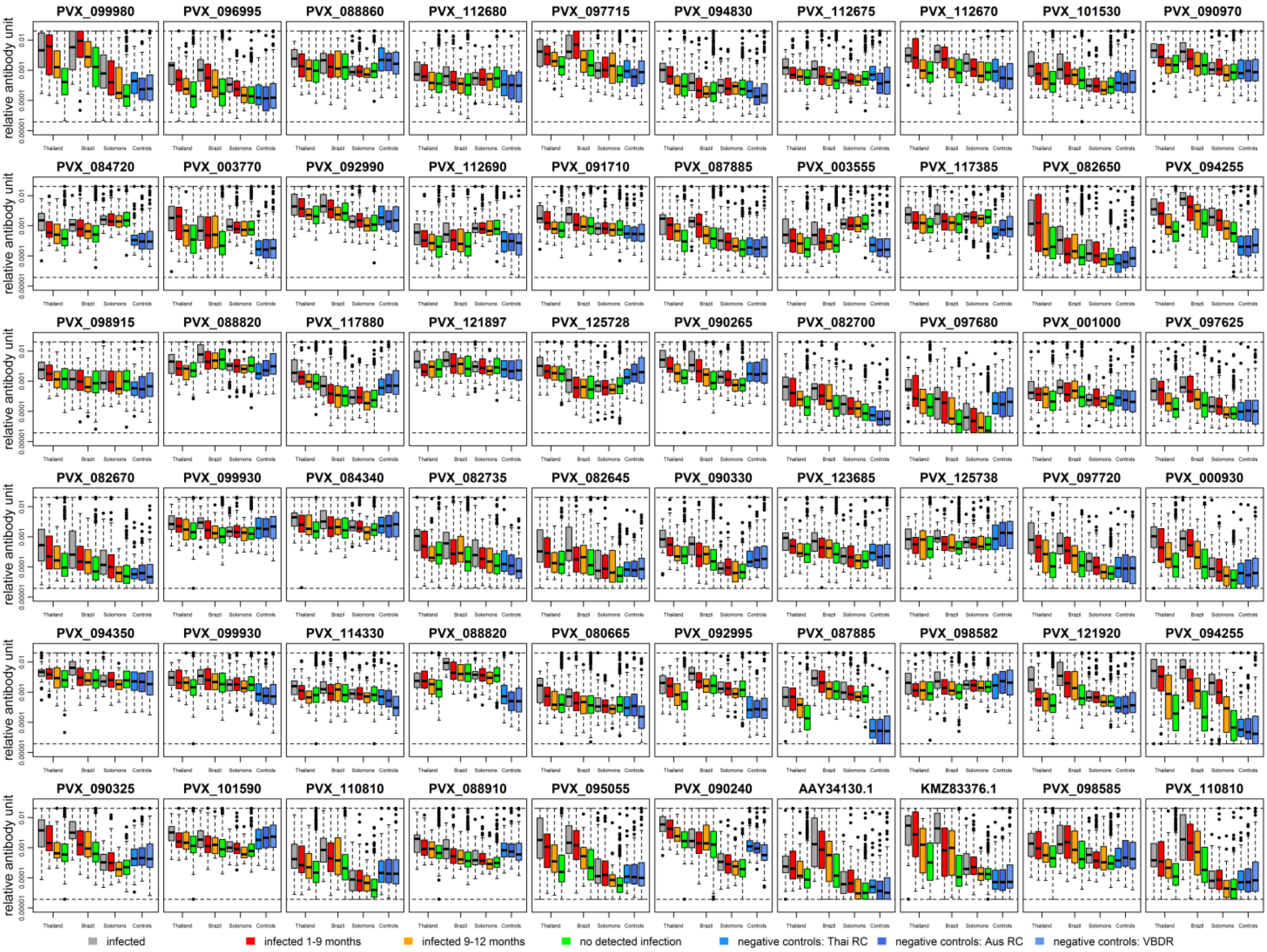


**Figure S5:** Measured antibody responses to 60 proteins on the Luminex® platform, stratified by geographical location and time since last PCR detected infection.

#### 2.2. Targets for classification

The data in Figure S5 demonstrate that for many proteins, there is a clear relationship between measured antibody response and time since last blood-stage *P. vivax* infection. Data from several proteins may be combined to optimise classification performance, however, first it is necessary to provide a definition of what we are attempting to classify and how optimal performance is selected.

***Why 9 months?*** In theory it is possible to provide quantitative estimates of the time since last infection, however a more useful outcome in practice is whether an individual has been infected within some past time period. Throughout this analysis, we have investigated the potential for measurements of antibody response from a sample to identify whether the sampled individual had a PCR-detected blood-stage *P. vivax* infection within the last 9 months. This time period was selected because:

(i) In the geographic regions studied, relapses are expected to occur at a frequency of every 1 – 2 months, with hypnozoites remaining in the liver for up to 1 – 2 years (Battle 2014, White 2016). Although some individuals are expected to be infected with liver-stage hypnozoites for greater than 1 year, the highest incidence of relapses is anticipated to occur within the first 9 months following mosquito bite infection (White 2011). Therefore, an individual with blood-stage *P. vivax* parasites detected by PCR within the last 9 months is expected to be a likely hypnozoite carrier.

(ii) When specifying a threshold for time since last infection, it is optimal if this threshold falls within a region where we have data. This allows us to have observed infections both before and after the threshold. The three longitudinal cohorts studied had follow-up for 12 months, thus 9 months falls within this window.

***What classification performance are we targeting?*** An ideal diagnostic tool would achieve 100% sensitivity (where all infected individuals are correctly identified as infected) and 100% specificity (where all non-infected individuals are correctly identified as non-infected). In reality there is a trade-off between sensitivity and specificity, which can be characterised by a Receiver

Operating Characteristic (ROC) curve. The particular balance between sensitivity and specificity will be selected according to the public health needs of the proposed diagnostic tool. Here we define three target product profiles (TPP) based on sensitivity-specificity trade-off (Ding 2017).

(i) Medium-sensitivity & medium-specificity (e.g. sens = 80% & spec = 80%). This target assigns equal weight to sensitivity and specificity. This target can be optimised by maximising the area under the ROC curve (AUC).
(ii) High-sensitivity & low-specificity (e.g. sens = 95% & spec = 50%). This target would be useful if one wishes to identify and treat all likely hypnozoite carriers, but some degree of over-treatment is acceptable.
(iii) Low-sensitivity & high-specificity (e.g. sens = 50% & spec = 95%). This target would be appropriate in surveillance settings for verifying the absence of transmission, where false-positives would incorrectly indicate ongoing transmission. The issue of false-negatives can be overcome by increasing survey size.

#### 2.3. Initial search of protein combinations

Here we investigate how combinations of the 60 proteins in Figure S5 were used to identify individuals with blood-stage *P. vivax* parasites detected by PCR within the last 9 months. A linear discriminant analysis (LDA) classification algorithm was used for this search strategy. There were 1,770 ways of choosing two out of 60 proteins, 34,220 ways of choosing three proteins, 487,635 ways of choosing four proteins, and 5,461,512 ways of choosing five proteins. All combinations up to size four were exhaustively evaluated to optimise classification performance for the three target product profiles (TPP) defined in section 2.2.

To investigate combinations of size five, we identified the 100 best combinations of four proteins for each of the three TPPs. We added all possible remaining proteins to each of these 300 combinations, and then assessed the classification performance of all of these combinations of size five. A similar procedure was implemented to investigate classification performance of combinations of proteins up to size eight. Figure S6 shows an overview of classification performance.

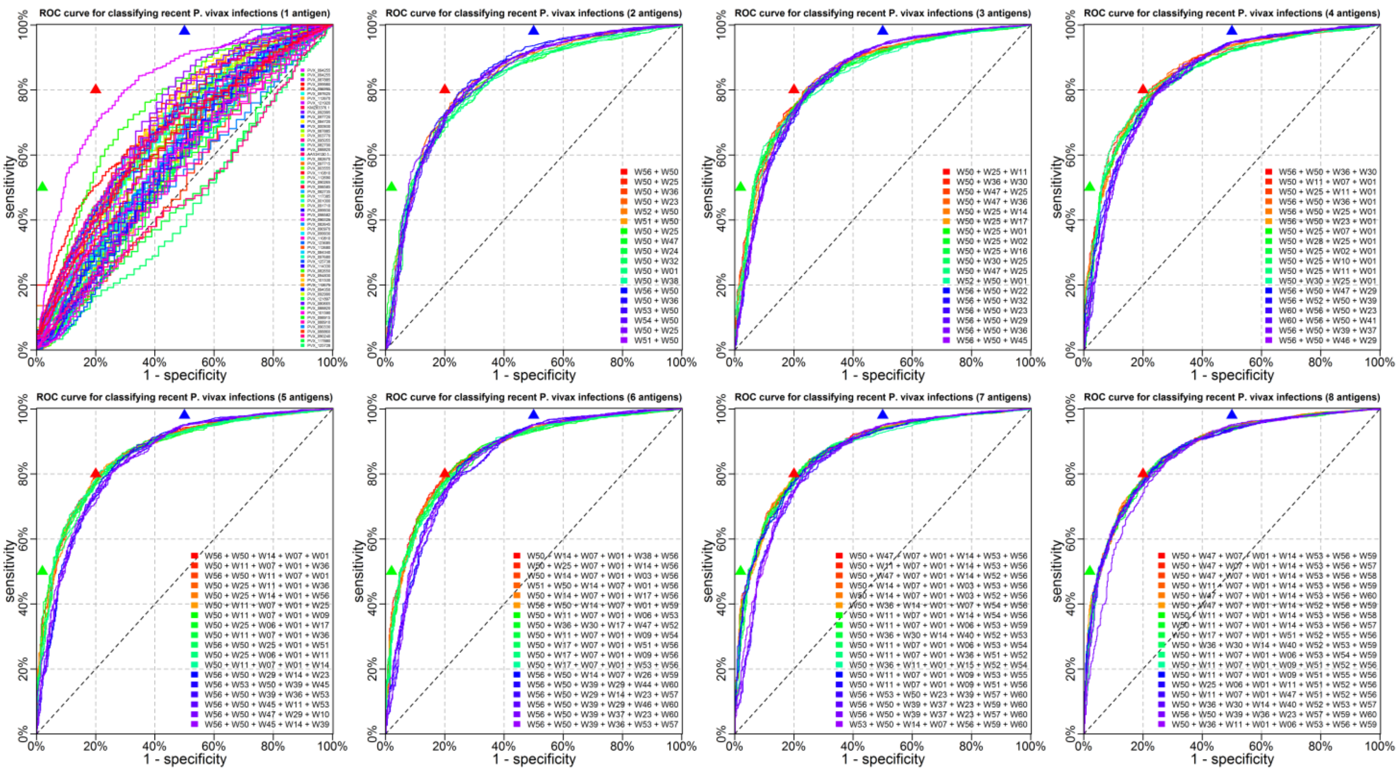


**Figure S6:** Performance of LDA classifier for combinations of proteins up to size eight for identifying individuals with blood-stage *P. vivax* infection within the last 9 months. The same training and testing data sets were used: data from endemic cohorts in Thailand, Brazil and the Solomon Islands plus negative control cohorts. The three triangles represent the TPPs in section 2.2. The red triangle is the medium-sensitivity & medium-specificity target, the blue triangle is the high-sensitivity & low-specificity target, and the green triangle is the low-sensitivity & high specificity target.

A network diagram was used to visualise which combinations of proteins provided the best classification performance (Figure S7). This network was generated based on the calculated AUC from ROC curves generated by an LDA classifier applied to all combinations of 4 from 60 proteins.

Denote $c_n$ where $1 \leq n \leq N = 487{,}635$ to be a combination of four proteins, and let $\text{AUC}_n$ be the area under the ROC curve generated by linear discriminant analysis. First re-order $n$ so that $\text{AUC}_n$ is decreasing. We now calculate a vector of vertex weights $V_j$ for each protein, and a matrix of edge weights $M_{jk}$ for connections between proteins. The vertex weights are proportional to how frequently a protein is selected in combinations with high ranking AUC.

$$V_j = \exp\left( -C_V \sum_{n=1}^{N} \mathbf{1}\left( j \in c_n \right) n \right) \qquad \text{(S11)}$$

The edge weights are proportional to how frequently pairs of proteins are selected in combinations with high ranking AUC.

$$M_{jk} = \exp\left( -C_M \sum_{n=1}^{N} \mathbf{1}\left( j \in c_n \right) \mathbf{1}\left( k \in c_n \right) n \right) \qquad \text{(S12)}$$

$C_v$ and $C_M$ are scaling constants manually selected to ensure good network visualisations. The network corresponding to the matrix $M_{jk}$ was plotted using the igraph R package with a Fruchterman-Reingold network layout. The sizes of the plotted vertices were scaled using $V_i$. The generated network of antigen combinations is shown in Figure S7.

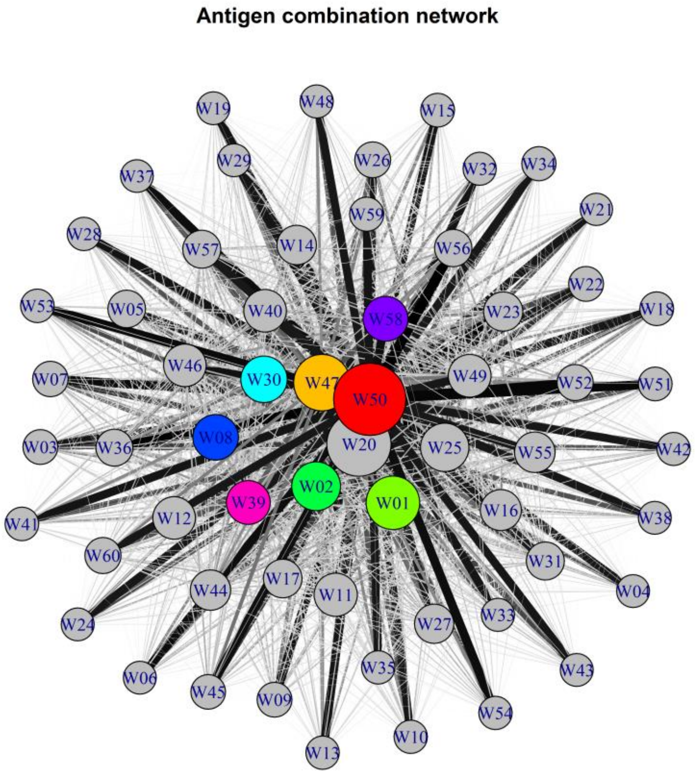


**Figure S7:** Network visualization of antigens selected in combinations of size 4. Larger vertices denote antigens frequently selected in combinations that maximise area under the curve (AUC) for classification. W20 was not selected because it was a construct of the same protein as W50 (PVX_094255).

##### 2.4. Cross-validated classification algorithms

Let $X_{ij}$ denote the matrix of measured log antibody titres in individual $i$ to protein $j$. Let $z_i$ denote whether individual $i$ was infected in the last 9 months ( $z_i = 1$ ) or not ( $z_i = 0$ ). The classification performance of algorithms is assessed through cross-validation whereby 2/3 of individuals are randomly selected to be in a training data set ($X_{ij,\text{train}}$ and $z_{i,\text{train}}$), and the remaining 1/3 selected in the testing data set ($X_{ij,\text{test}}$ and $z_{i,\text{test}}$). This process is repeated 500 times.

###### 2.4.1. Statistical classifiers

A number of conventional classifiers from the fields of statistical inference and machine learning were repeatedly applied to the training and testing data sets. Classifiers considered were:

(i) logistic regression
(ii) LDA
(iii) quadratic discriminant analysis (QDA)
(iv) decision trees
(v) random forests

These classifiers are widely used and are described in detail by Hastie, Tibshirani & Friedman. All algorithms were implemented in R. Decision trees were implemented using the rpart R package. Random forests were implemented using the randomForest R package.

The random forests algorithm can also be used to generate a variable importance plot which can provide a ranking of antigens in terms of their contribution to classification performance (Figure S8).

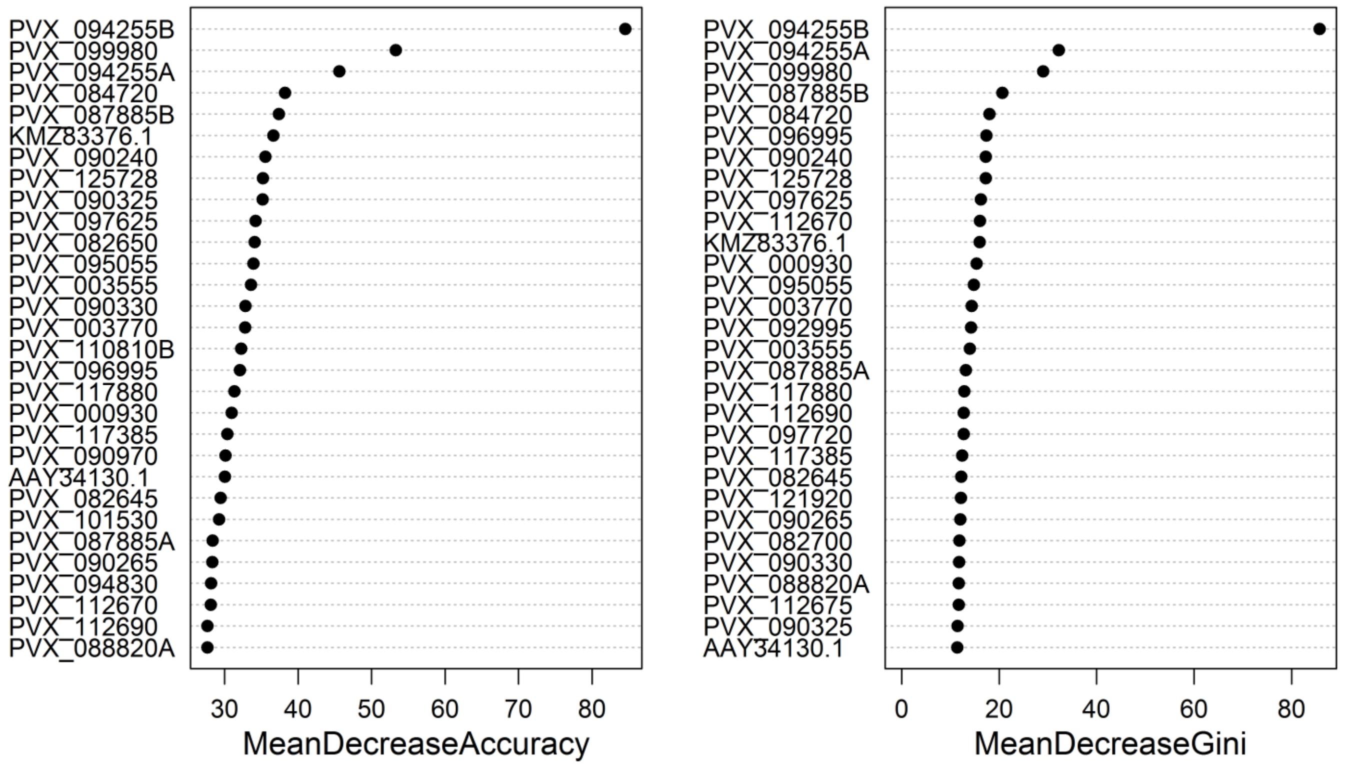


**Figure S8:** Variable importance plot for identifying antigens that contribute to optimal classification in a random forests algorithm generated using the varImpPlot function from the randomForest R package. Note the high degree of concordance between the listed proteins and those in the network in Figure S7.

##### 2.4.2. Antibody dynamics classifiers

Section 2.4.1 lists the 'off-the-shelf' statistical classifiers. Here we describe how information on antibody dynamics can be incorporated into a classification algorithm.

***Antibody dynamics algorithm #1***

This is similar to the LDA and QDA algorithms. Let $\alpha_{\text{train,0-9m}}$ and $\Sigma_{\text{train,0-9m}}$ be the mean and covariance matrix of the multi-variate normal distribution fitted to data from recently infected individuals in the training data. Let $\alpha_{\text{train,9m+}}$ and $\Sigma_{\text{train,9m+}}$ be the mean and covariance matrix of the multi-variate normal distribution fitted to data from individuals without recent infection in the training data. For a measurement of antibody responses from an individual in the testing data set $x_{i,\text{test}}$ we can calculate the relative probabilities as follows:

$$\begin{aligned} Q\left(z_{i,\text{test}}=0\middle|x_{i,\text{test}}\right) &= \frac{1}{\sqrt{2\pi\left|\Sigma_{\text{train,9m+}}\right|}} e^{-\frac{1}{2}\left(\left(x_{i,\text{test}}-\alpha_{\text{train,9m+}}\right)^T \Sigma_{\text{train,9m+}}^{-1}\left(x_{i,\text{test}}-\alpha_{\text{train,9m+}}\right)\right)} \\ Q\left(z_{i,\text{test}}=1\middle|x_{i,\text{test}}\right) &= \frac{1}{\sqrt{2\pi\left|\Sigma_{\text{train,0-9m}}\right|}} e^{-\frac{1}{2}\left(\left(x_{i,\text{test}}-\alpha_{\text{train,0-9m}}\right)^T \Sigma_{\text{train,0-9m}}^{-1}\left(x_{i,\text{test}}-\alpha_{\text{train,0-9m}}\right)\right)} \end{aligned} \tag{S13}$$

These probabilities can be normalised as follows:

$$\begin{aligned} P\left(z_{i,\text{test}}=0\middle|x_{i,\text{test}}\right) &= \frac{Q\left(z_{i,\text{test}}=0\middle|x_{i,\text{test}}\right)}{Q\left(z_{i,\text{test}}=0\middle|x_{i,\text{test}}\right)+Q\left(z_{i,\text{test}}=1\middle|x_{i,\text{test}}\right)} \\ P\left(z_{i,\text{test}}=1\middle|x_{i,\text{test}}\right) &= \frac{Q\left(z_{i,\text{test}}=1\middle|x_{i,\text{test}}\right)}{Q\left(z_{i,\text{test}}=0\middle|x_{i,\text{test}}\right)+Q\left(z_{i,\text{test}}=1\middle|x_{i,\text{test}}\right)} \end{aligned} \tag{S14}$$

***Antibody dynamics algorithm #2***

This algorithm extends antibody dynamics algorithm #1 by accounting for the data available in the known time since last infection in the training data set $T_{i,\text{train}}$. The first step is to use a linear regression model on those individuals with a detected blood-stage infection to estimate the geometric mean antibody titre at the time of last PCR-detectable blood-stage infection and the decay rate of antibodies over time:

$$X_{ij,\text{train}} \sim \log\left(\alpha_j^0\right) + r_j^0 T_{i,\text{train}} + \varepsilon_j \qquad \text{(S15)}$$

where $X_{ij,\text{train}}$ is the log antibody titer to protein $j$ from individual $i$ in the training data set, $T_{i,\text{train}}$ is the time since last PCR-detected blood-stage infection in individual $i$ in the training data set, $r_j^0$ is the estimated antibody decay rate, log( $\alpha_j^0$ ) is the intercept term, i.e. the log of the geometric mean titre at the time of infection, and $\varepsilon_j$ is Normally distributed measurement error. Note that in contrast to equation (S1), the serological data utilised here are from a cross-sectional and not a longitudinal survey, therefore the resulting regression model does not account for mixed effects.

We can now follow the approach outlined above in Section 1.3.

$$\begin{aligned}\mathbf{P}\left(x_i \,|\, t, z_i = 0\right) &= \left(2\pi\right)^{-\frac{J}{2}} \left|\Sigma_{\text{train,9m+}} + t^2\Sigma_r\right|^{-\frac{1}{2}} e^{-\frac{1}{2}\left(x_i - \alpha^0 - r^0 t\right)^T \left(\Sigma_{\text{train,9m+}} + t^2\Sigma_r\right)^{-1}\left(x_i - \alpha^0 - r^0 t\right)} \\ \mathbf{P}\left(x_i \,|\, t, z_i = 1\right) &= \left(2\pi\right)^{-\frac{J}{2}} \left|\Sigma_{\text{train,0-9m}} + t^2\Sigma_r\right|^{-\frac{1}{2}} e^{-\frac{1}{2}\left(x_i - \alpha^0 - r^0 t\right)^T \left(\Sigma_{\text{train,0-9m}} + t^2\Sigma_r\right)^{-1}\left(x_i - \alpha^0 - r^0 t\right)}\end{aligned} \qquad \text{(S16)}$$

And these can be inverted similar to above to give

$$\begin{aligned}\mathbf{P}\left(z_i = 1 \middle| x_i\right) &= \int_0^9 \mathbf{P}\left(x_i \,|\, t, z_i = 1\right) dt \\ \mathbf{P}\left(z_i = 0 \middle| x_i\right) &= \int_9^{t_{\max}} \mathbf{P}\left(x_i \,|\, t, z_i = 0\right) dt\end{aligned} \qquad \text{(S17)}$$

##### 2.4.3. Composite classification algorithm

Figure S9 shows a comparison of the cross-validated classification performance of the seven algorithms described above on several data sets. All algorithms provide comparable performance, indicating that they all capture the same signal in the data. For the combination of eight proteins, the Random Forests algorithm performs the best on average. The antibody dynamics algorithm 2 performs best at hitting high specificity targets. These seven algorithms can be designated as components of a composite classification algorithm. For a desired target of sensitivity and specificity, the composite classification algorithm outputs the results of the best performing component algorithm.

The same concept can be applied to combinations of proteins. For example, there are 28 ways to choose a combination of two proteins from eight proteins. Thus we can generate 28 ROC curves and choose the one that performs best for a given target of sensitivity and specificity.

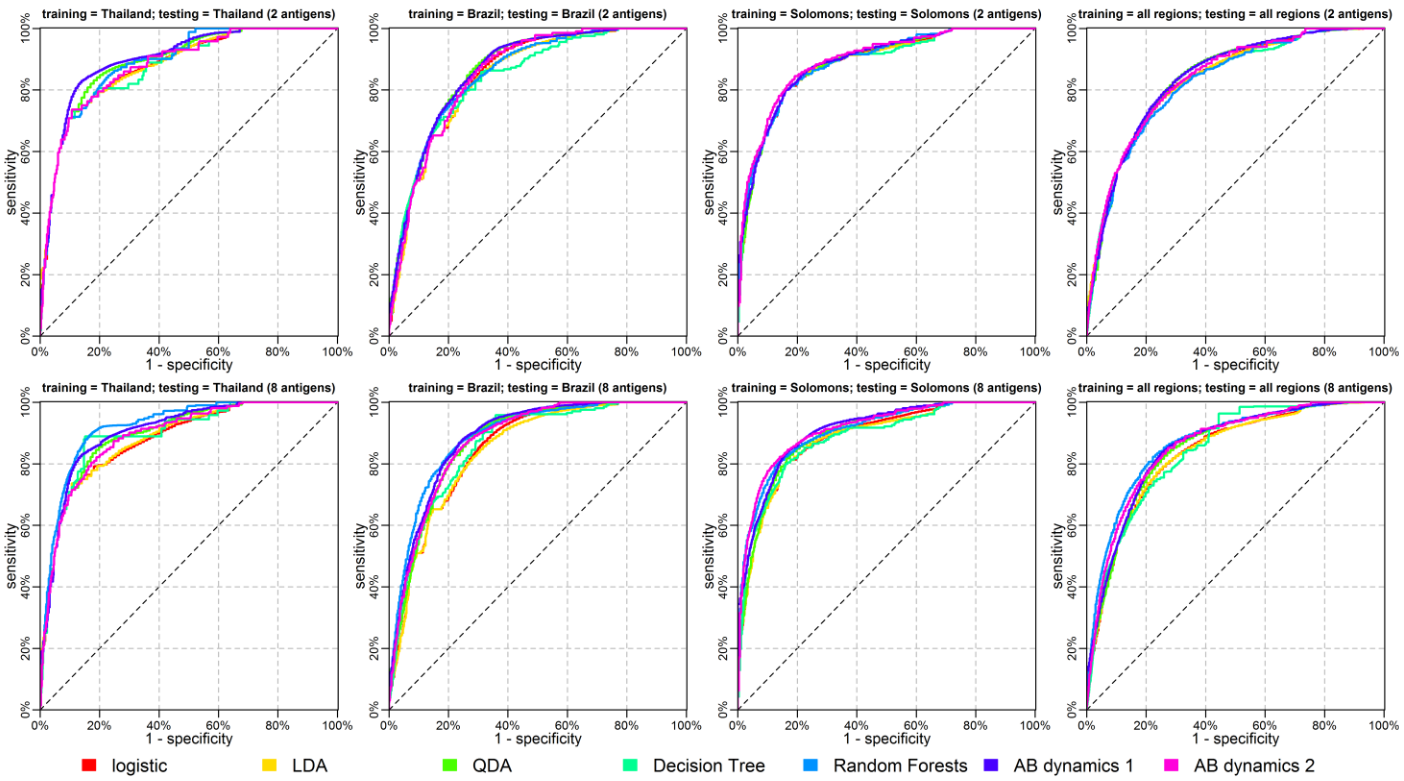


**Figure S9:** Receiver operating characteristic (ROC) curves depicting comparison of cross-validated classification performance for the seven classification algorithms considered. For a given sensitivity and specificity target, the composite classification algorithm selects the best of the seven component algorithms. Its ROC curve can be considered as the convex hull of the ROC curves for the seven component algorithms. All curves presented are the median of 500 repeat cross-validations.

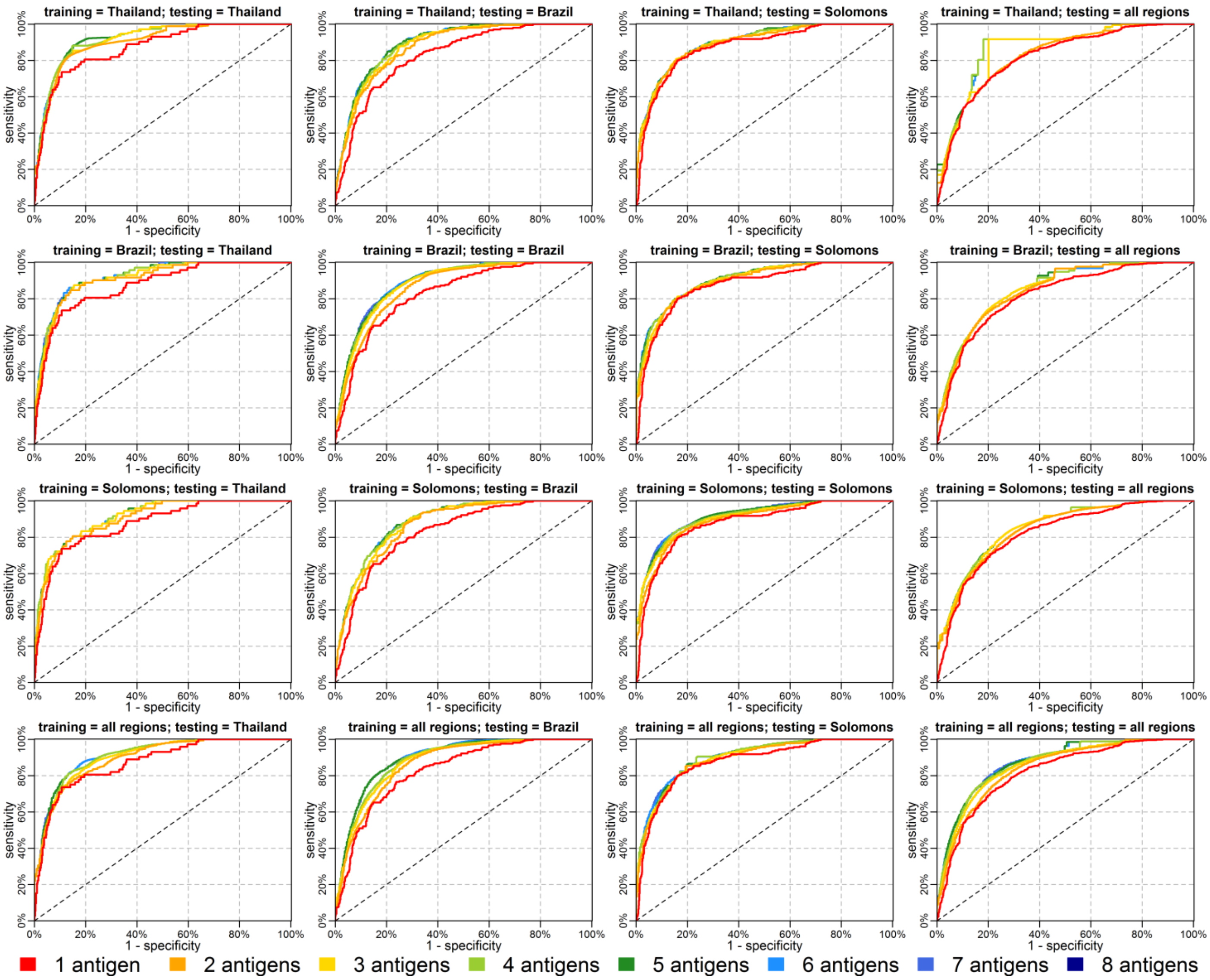


**Figure S10:** Receiver operating characteristic (ROC) curve for the composite classification algorithm. All curves presented are the median of 500 repeat cross-validations.

In Figure S10 we see that the composite classification algorithm has comparable performance across a wide range of training and testing data sets. For example, when trained on Brazilian data, the classifier performs well on Thai data. This repeated pattern suggests that an algorithm trained in one epidemiological setting can be applied to other epidemiological settings.

#### 2.5. Supplementary results

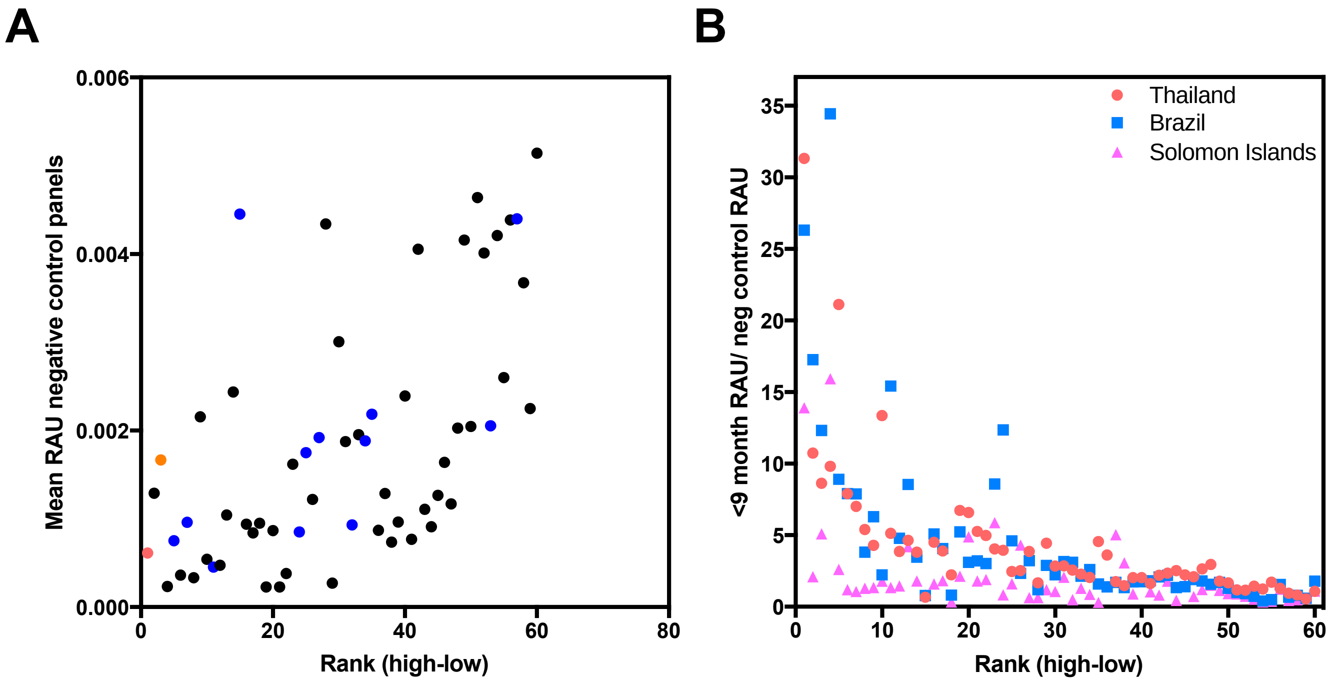


**Figure S11:** Association between background reactivity in non-malaria exposed controls and ranking of candidate SEMs. A) Mean relative antibody units (RAU) detected in malaria-naïve control panels from Melbourne, Australia (n=202) and Bangkok, Thailand (n=72) compared to the ranking of the 60 candidate *P.vivax* proteins generated during the validation phase. WGCF expressed proteins are in black and *E. coli* or Baculovirus expressed proteins are in blue. RBP2b$_{161\text{-}1454}$ (*E. coli*) is in red and RBP2b$_{1986\text{-}2653}$ is in orange. In part B) the ranking has been compared to the ratio of the antibody level detected in individuals with a recent *P.vivax* infection (previous 9 months) compared to that detected in the malaria-naïve controls.

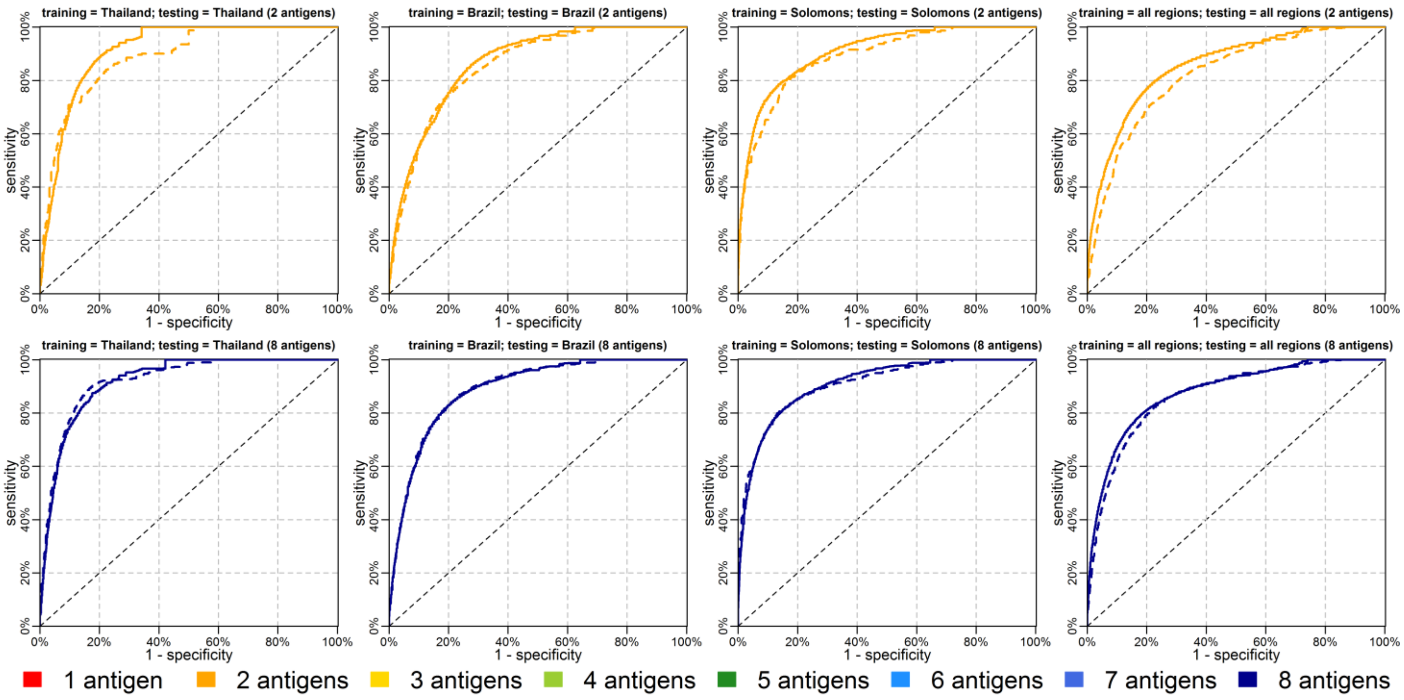


**Figure S12:** Assessment of incorporating information on an individual's age into a Random Forests classifier. Age was incorporated as a continuous variable in years. The Random Forests classifier was selected for this sensitivity analysis because of the flexibility with which it could incorporate information on age. Dashed lines denote predictions without age, and solid lines denote predictions with age.

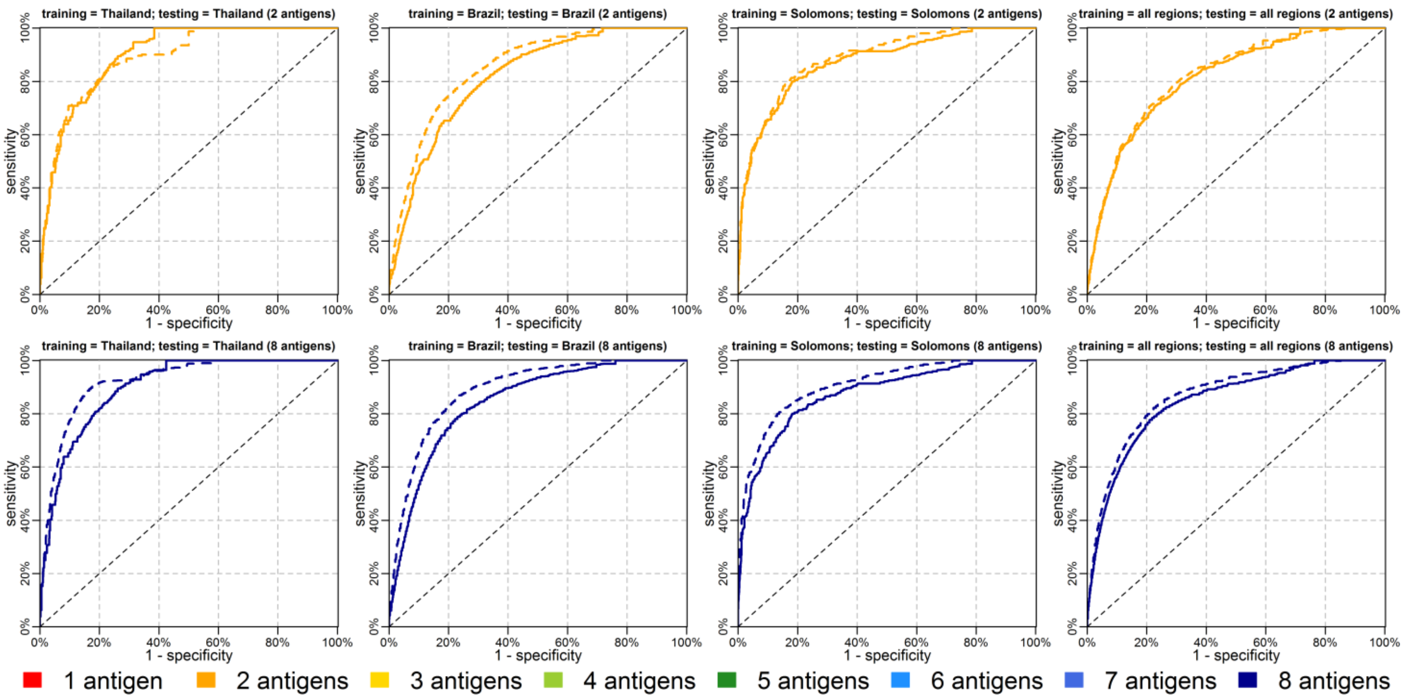


**Figure S13:** Assessment of the role of removing data from negative control participants on classification performance using a Random Forests classifier. Dashed lines denote predictions with negative control participants included, and solid lines denote predictions with negative control participants excluded.

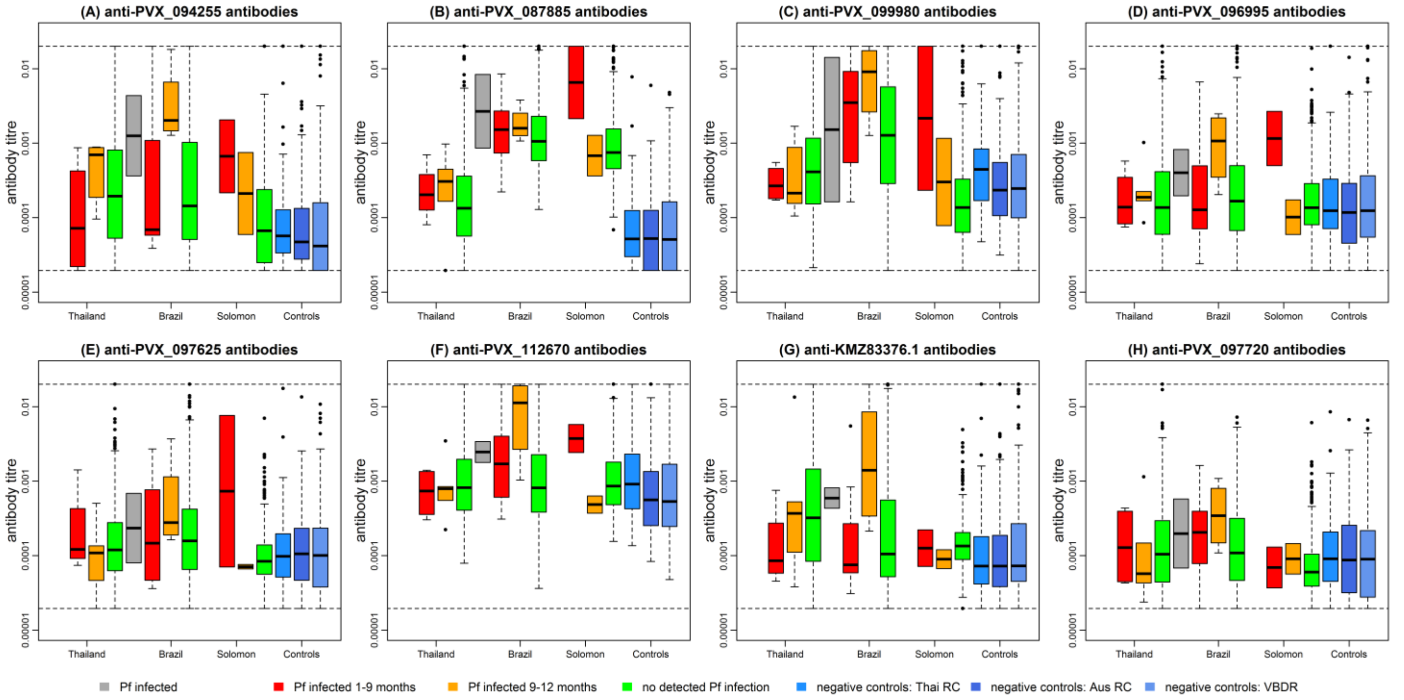


**Figure S14:** Association between measurements of our top eight *P. vivax* proteins and time since last PCR detected blood-stage *P. falciparum* infection. Note that individuals with any sample PCR positive for *P. vivax* have been removed. For all eight proteins in each of the three cohorts, there was no significant difference in relative antibody units between individuals with no PCR-detected blood-stage *P. falciparum* and individuals with a blood-stage *P. falciparum* in the last 9 months (two-sided Student's t-test).
