## Supplementary material for "Development and validation of serological markers for detecting recent exposure to *Plasmodium vivax* infection"

**Table S2.** Purified *P. vivax* proteins used in the validation phase and their individual performance, complete list. Area under the curve (AUC) from the single antigen classifier is shown; proteins are listed in order of best performance. All 60 proteins are shown; Table 2 in the main manuscript included the first 8 proteins. References listed are for the protein production and purification method.

| Protein ID^a^ | Gene Annotation^a^ | Short code | Protein length, aa | Construct, aa (size) | Expression System^c^ | Purification Method | AUC |
| --- | --- | --- | --- | --- | --- | --- | --- |
| PVX_094255B | reticulocyte binding protein 2b (RBP2b) | W50 | 2806 | 161-1454 (1294) | *E. coli* | 2x affinity + size exclusion (*1*) | 0.819 |
| PVX_099980 | merozoite surface protein 1 (MSP1-19) | W01 | 1751 | 1622-1729 (108) | WGCF | One-step Ni column | 0.804 |
| PVX_000930 | sexual stage antigen s16, putative | W40 | 140 | 31-end (110) | WGCF | One-step Ni column | 0.78 |
| PVX_094255A | reticulocyte binding protein 2b (RBP2b) | W20 | 2806 | 1986-2653 (667) | WGCF | One-step Ni column | 0.779 |
| PVX_087885B | rhoptry-associated membrane antigen, putative | W47 | 730 | 462-730 (269) | WGCF | One-step Ni column (*2*) | 0.774 |
| PVX_095055 | Rh5 interacting protein, putative (RIPR) | W55 | 1075 | 552-1075 (524) | *E. coli* | 2x affinity + size exclusion (*3*) | 0.767 |
| AAY34130.1^b^ | Duffy binding protein (DBP, region 2, AH strain) | W57 | 237 | 1-237 (237) | *E. coli* | Ni, ion exchange, gel filtration (*4*) | 0.764 |
| PVX_097715 | hypothetical protein | W05 | 450 | 20-end (431) | WGCF | One-step Ni column | 0.76 |
| PVX_110810B | Duffy binding protein (DBP, region 3-5, Sal1 strain) | W60 | 1070 | 508-899 (392) | *E. coli* | One-step Ni column (*3*) | 0.757 |
| PVX_097625 | merozoite surface protein 8 (MSP8), putative | W30 | 487 | 24-463 (440) | WGCF | One-step Ni column | 0.756 |
| PVX_087885A | rhoptry associated membrane antigen, putative | W16 | 730 | 462-730 (269) | WGCF | One-step Ni column | 0.751 |
| PVX_092995 | tryptophan-rich antigen (Pv-fam-a) | W46 | 385 | 25-358 (334) | WGCF | One-step Ni column (*2*) | 0.748 |
| PVX_097720 | merozoite surface protein 3 (MSP3.10) | W39 | 852 | 25-end (828) | WGCF | One-step Ni column | 0.74 |
| PVX_112670 | unspecified product | W08 | 335 | 34-end (302) | WGCF | One-step Ni column | 0.738 |
| PVX_096995 | tryptophan-rich antigen (Pv-fam-a) | W02 | 480 | 61-end (420) | WGCF | One-step Ni column | 0.733 |
| PVX_121920 | reticulocyte binding protein 2a (RBP2a) | W49 | 2487 | 160-1135 (976) | *E. coli* | 2x affinity + size exclusion (*1*) | 0.731 |
| PVX_082700 | merozoite surface protein 7 (MSP7.1) | W27 | 420 | 23-end (397) | WGCF | One-step Ni column | 0.729 |
| KMZ83376.1^b^ | erythrocyte binding protein II (PvEBPII) | W58 | 786 | 109-432 (324) | *E. coli* | Ni, ion exchange, gel filtration (*4, 5*) | 0.725 |
| PVX_090325 | reticulocyte binding protein 2c (RBP2c non binding region) | W51 | 2824 | 501-1300 (800) | *E. coli* | 2x affinity + size exclusion (*1*) | 0.709 |
| PVX_088820B | tryptophan-rich antigen (Pv-fam-a) | W44 | 316 | 58-end (259) | WGCF | One-step Ni column (*2*) | 0.701 |
| PVX_084720 | translocon component PTEX150, putative | W11 | 908 | 24-908 (885) | WGCF | One-step Ni column | 0.688 |
| PVX_082670 | merozoite surface protein 7 (MSP7), putative | W27 | 411 | 24-end (388) | WGCF | One-step Ni column | 0.683 |
| PVX_110810A | Duffy binding protein (DBP, region 2, Sal1 strain) | W53 | 1070 | 193-521 (329) | *E. coli* | Ni, ion exchange, gel filtration (*4, 5*) | 0.681 |
| PVX_090265 | tryptophan-rich antigen (Pv-fam-a) | W26 | 326 | 1-326 (326) | WGCF | One-step Ni column | 0.68 |
| PVX_098585 | reticulocyte binding protein 1a (RBP1a) | W59 | 2833 | 160-1170 (1011) | *E. coli* | 2x affinity + size exclusion (*1*) | 0.675 |
| PVX_003770 | merozoite surface protein 5 (MSP5) | W12 | 387 | 23-365 (343) | WGCF | One-step Ni column | 0.669 |
| PVX_091710 | hypothetical protein, conserved | W15 | 1689 | 26-884 (859) | WGCF | One-step Ni column | 0.658 |
| PVX_099930B | high molecular weight rhoptry protein 2 (RhopH2) | W42 | 1369 | 23-387 (365) | WGCF | One-step Ni column (*2*) | 0.65 |
| PVX_090970 | hypothetical protein, conserved | W10 | 266 | 20-254 (235) | WGCF | One-step Ni column | 0.645 |
| PVX_123685 | histone-lysine N-methyltransferase, H3 lysine-4 specific (SET10), putative | W37 | 1963 | 1320-end (644) | WGCF | One-step Ni column | 0.639 |
| PVX_101530 | Plasmodium exported protein, unknown function | W09 | 367 | 38-end (330) | WGCF | One-step Ni column | 0.637 |
| PVX_082735 | thrombospondin-related anonymous protein (TRAP) | W34 | 556 | 26-493 (468) | WGCF | One-step Ni column | 0.636 |
| PVX_082645 | merozoite surface protein 7 (MSP7), putative | W35 | 377 | 23-end (355) | WGCF | One-step Ni column | 0.636 |
| PVX_092990 | tryptophan-rich antigen (Pv-fam-a) | W13 | 1414 | 1126-1414 (289) | WGCF | One-step Ni column | 0.635 |
| PVX_097680 | merozoite surface protein 3 (MSP3.3) | W28 | 1016 | 21-end (996) | WGCF | One-step Ni column | 0.635 |
| PVX_001000 | hypothetical protein, conserved | W29 | 668 | 20-end (650) | WGCF | One-step Ni column | 0.633 |
| PVX_112690 | unspecified product | W14 | 313 | 30-313 (284) | WGCF | One-step Ni column | 0.625 |
| PVX_003555 | Plasmodium exported protein, unknown function | W17 | 1122 | 434-1075 (642) | WGCF | One-step Ni column | 0.624 |
| PVX_090330 | reticulocyte binding protein 2 precursor (PvRBP-2), putative | W36 | 623 | 31-141 (111) | WGCF | One-step Ni column | 0.614 |
| PVX_117385 | phosphatidylinositol-4-phosphate-5-kinase, putative | W18 | 326 | 1-326 (326) | WGCF | One-step Ni column | 0.612 |
| PVX_099930A | high molecular weight rhoptry protein 2 (RhopH2) | W32 | 1369 | 23-387 (365) | WGCF | One-step Ni column | 0.611 |
| PVX_094350 | hypothetical protein, conserved | W41 | 1220 | 2-615 (614) | WGCF | One-step Ni column (*2*) | 0.609 |
| PVX_080665 | hypothetical protein, conserved | W45 | 553 | 25-553 (529) | WGCF | One-step Ni column (*2*) | 0.603 |
| PVX_114330 | Plasmodium falciparum CPW-WPC domain containing protein | W43 | 185 | 24-185 (162) | WGCF | One-step Ni column (*2*) | 0.602 |
| PVX_112675 | unspecified product | W07 | 312 | 33-end (280) | WGCF | One-step Ni column | 0.601 |
| PVX_112680 | unspecified product | W04 | 313 | 33-end (281) | WGCF | One-step Ni column | 0.6 |
| PVX_082650 | merozoite surface protein 7 (MSP7), putative | W19 | 453 | 24-end (429) | WGCF | One-step Ni column | 0.593 |
| PVX_094830 | hypothetical protein, conserved | W06 | 250 | 19-end (232) | WGCF | One-step Ni column | 0.588 |
| PVX_098582 | reticulocyte binding protein 1b (RBP1b) | W48 | 2608 | 140-1275 (1136) | *E. coli* | 2x affinity + size exclusion (*1*) | 0.587 |
| PVX_088820A | tryptophan-rich antigen (Pv-fam-a) | W22 | 316 | 58-end (259) | WGCF | One-step Ni column | 0.585 |
| PVX_084340 | IMP-specific 5'-nucleotidase, putative | W33 | 444 | 1-444 (444) | WGCF | One-step Ni column | 0.582 |
| PVX_090240 | cysteine-rich protective antigen, putative (CyRPA) | W56 | 366 | 27-366 (340) | Baculovirus | 1x affinity + size exclusion (*5*) | 0.58 |
| PVX_121897 | tryptophan-rich antigen (Pv-fam-a) | W24 | 275 | 24-end (252) | WGCF | One-step Ni column | 0.579 |
| PVX_125738 | unspecified product | W38 | 786 | 1-786 (786) | WGCF | One-step Ni column | 0.577 |
| PVX_098915 | subpellicular microtubule protein 1 (SPM1), putative | W21 | 521 | 1-521 (521) | WGCF | One-step Ni column | 0.563 |
| PVX_088910 | GPI-anchored micronemal antigen, putative (GAMA) | W54 | 771 | 22-551 (530) | *E. coli* | Two-step affinity (*3*) | 0.563 |
| PVX_101590 | reticulocyte-binding protein 2 (RBP2), like (RBP2-P2) | W52 | 641 | 161-641 (481) | *E. coli* | 2x affinity + size exclusion (*5*) | 0.561 |
| PVX_088860 | sporozoite invasion-associated protein 2 (SIAP2), putative | W03 | 412 | 33-end (380) | WGCF | One-step Ni column | 0.55 |
| PVX_125728 | unspecified product | W25 | 279 | 30-end (250) | WGCF | One-step Ni column | 0.544 |
| PVX_117880 | rhoptry neck protein 2 (RON2), putative | W23 | 2203 | 21-198 (178) | WGCF | One-step Ni column | 0.504 |
| ^a^PlasmoDB release 36 (http://plasmodb.org/plasmo/), ^b^GenBank | |  |  |  |  |  |  |
